# Evaluating selection at intermediate scales within genes provides robust identification of genes under positive selection in *M. tuberculosis* clinical isolates

**DOI:** 10.1101/2025.05.07.652684

**Authors:** Thomas R. Ioerger, Anthony Shatby

## Abstract

Multiple studies have reported genes in the *M. tuberculosis* (Mtb) genome that are under diversifying selection, based on genetic variants among Mtb clinical isolates. These might reflect adaptions to selection pressures associated with modern clinical treatment of TB. Many, but not all, of these genes under selection are related to drug resistance. Most of these studies have evaluated selection at the gene-level. However, positive selection can be evaluated on different scales, including individual sites (codons) and local regions within an ORF. In this paper, we use GenomegaMap, a Bayesian method for calculating selection, to evaluate selection of genes in the Mtb genome at all three levels. We present evidence that the intermediate analysis (windows of codons) yields the most credible list of candidate genes under selection (excluding PPE and PE_PGRS genes, which are predicted less reliably due to frequent sequencing errors). A further advantage of this approach is that it identifies specific regions within proteins that are under selective pressure, which is useful for structural and functional interpretation. In an analysis of two separate collections of Mtb clinical isolates (from Moldova; and a globally-representative set), we observed 53 and 173 significant genes under selection, with 36% overlap. The lists of genes under selection include many drug-resistance genes, as well as other genes that have previously been reported to be under selection (*resR, phoR*). The specific regions under selection identified within drug-resistance genes are shown to correspond to protein structural features known to be involved in resistance, supporting accuracy of the method. Positive selection in several ESX-1-related genes was also observed, suggesting adaptation to immune pressure.

## 1. INTRODUCTION

Recent studies examining large collections of genome sequences of *M. tuberculosis* (Mtb) clinical isolates have reported the identification of genes that appear to be under positive selection [1–3]. Traditionally, positive selection is indicated by an excess of non-synonymous over synonymous mutations (in protein-coding regions), although clustering of SNPs (e.g. in non-coding regions), homoplasy [4], and other measures of convergent evolution [5] have also been used in the literature. This is taken to imply on-going evolution and adaptation to modern chemotherapy, including pressures due to antibiotic exposure, as well as adaptive mutations.

This is evidenced by the fact that genes associated with drug resistance are often found to have among the highest selection signals [6]. For example, genes directly involved as drug targets or activators (like *rpoB* or *katG*) exhibit selection, as well as genes with indirect effects, like efflux pumps (*tap, mmr, mmpL5*) and regulators (*whiB6, resR, phoR*) that might confer tolerance through changes in cell wall permeability, metabolism, etc. In addition, selection signals sometimes appear in genes due to compensatory mechanisms, where mutations in one gene help reduce the fitness costs of mutations in another [7].

Evaluating selection in ORFs (open reading frames) is usually done within a Bayesian framework, where observed mutations are used to infer posterior probability distributions over mutation rates that would best explain the observations. dN/dS is not just the ratio of raw counts of non-synonymous (NS) to synonymous (S) mutations, but is corrected for a) relative number of possibilities of NS and S changes for each codon (typically twice as many NS as S), and b) for evolutionary time [8]. Positive selection is defined as dN/dS>1. First, a null model is postulated where mutations occur randomly throughout an ORF and accumulate (exponentially) over time. Models such as NY98 [9] use a parameterized codon substitution matrix for rates of substitution between each pair of aligned codons, i and j, that differentiates transition vs transversion rates (using a latent variable k), and synonymous vs non-synonymous changes (using a latent variable omega, w), relative to background codon frequencies p_j_:

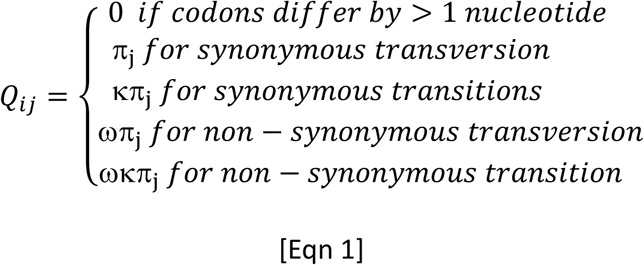

Over an epoch of evolutionary time *t*, the substitution matrix relaxes by the following equation: *P*(*t*) = *e*^*Qt*^. The likelihood function for observed allele frequencies, *f_i_*, is assumed to be *Multinomial(f_i_; P(t))*. Integrating out *t*, the marginal distribution for allele frequencies among extant sequences can be shown to follow a multinomial-Dirichlet distribution [10]. Observed mutations are mapped onto a phylogenetic tree to identify unique evolutionary events, and then the relative number of synonymous and non-synonymous events is used to compute posterior distributions over the parameters (including w) under different assumptions (such as whether there is a constant molecular clock, or rates can differ independently in each branch).

PAML [11] is a widely-used tool for evaluating selection in sequence data, which uses maximum-likelihood (ML) to estimate the model parameters. However, PAML does not scale-up well, and is difficult to run for collections of thousands of genomes, preventing its use in studies such as [1]. The complexity is derived from the requirement of large-scale phylogenetic trees for the computations. Several alternative approaches for evaluating selection have been proposed that are more efficient, enabling their use in analysis of large-scale collections. One such tool is GenomegaMap (https://github.com/danny-wilson/genomegaMap) [10]. GenomegaMap does not require a phylogenetic tree, aggregating SNPs without regard for when the mutations occurred and in which branches, but otherwise relies on a very similar Bayesian framework [Eqn 1] to explain the relative proportion of NS to S mutations in an ORF. GenomegaMap uses Markov Chain Monte Carlo (MCMC) methods (described in more detail below) to infer selection parameters for mutations observed in genes. GenomegaMap was in fact applied to analyze >10,000 Mtb genome sequences from the CRyPTIC collection (demonstrating its scalability) and proposed a list of genes under positive selection, which included many drug-resistance-related genes (consistent with expectations).

As with PAML, GenomegaMap implements several different models for evaluating selection at different levels of granularity. In addition to the gene-level analysis (referred to as the Constant model in GenomegaMap), where all mutations in an ORF are aggregated, selection can also be evaluated at the level of individual codons (referred to as the Independent model). Importantly, these two models have different sensitivities for detecting selection. While analysis of selection in individual codons might seem to have the highest resolution (signal-to-noise ratio), this analysis suffers from the inability to exploit information at multiple nearby sites in a region (such as in an active site, or protein-protein interaction site). On the other hand, while aggregating SNPs at the gene-level can take advantage by combining mutations at multiple sites within a gene, the signal for selection can be diluted by other unrelated mutations occurring in other less-conserved parts of the protein.

An intermediate approach would be to evaluate selection signals in localized regions throughout an ORF. Mutations that can similarly impact the function of a protein often involve multiple residues that cluster together in sequence (and hence in 3D structure), such as residues in an active site or a site of protein-protein interaction. This then produces the appearance of clusters of non-synonymous mutations in multiple alignments of genes under selection. To evaluate selection in such clustered regions, GenomegaMap offers a third mode, called the “Window” model, in which an ORF is divided into several contiguous segments, and an independent w parameter is used to model selection in each region. This model assumes selection pressure will be constant among codons within each region, but can differ between regions within an ORF. Importantly, the number and boundaries of these regions are themselves parameters of the model, and can be adjusted dynamically to maximize fit to mutation data. The advantage of this Window approach is that it produces a “smoothed” selection signal (over adjacent codons) by identifying regions within the ORF where there is a cluster of SNPs with a localized over-representation of non-synonymous to synonymous SNPs.

GenomegaMap uses MCMC sampling to simulate posterior probability distributions over all model parameters by generating a sequence of parameter combinations from a Markov chain; in the end, the mean value and credible interval for each parameter over the trajectory can be extracted (after removing an initial “burn-in” portion of the trajectory).

Comparing the quality of model fits based on total likelihood, Wilson [10] concluded that the codon-level analysis (Independent model, identifying genes with individual codons under positive selection) fit the data better than the Window model. However, using a criterion of Pr(w>1)>0.9 (dN/dS>1 in >90% of MCMC samples), thousands of genes were found to have at least one codon under selection (see Fig 3 in Wilson). While many drug-resistance-related genes were included, when genes were ranked by the proportion of codons under selection, some expected genes, like *gyrA, rpoB,* and *embB*, were found to have ranks in the range of 2000-3000 (out of ∼4000 genes). For the purposes of prioritization, it would be expected that such genes would be ranked closer to the top.

In this paper, we empirically compare the three different models of selection – at the gene-level, codon-level, and in local windows (gene segments) – by evaluating them on analysis of a set of ∼2000 Mtb clinical isolates from Moldova [12]. While thousands of genes have w>1 (as a point estimate), determining which genes are statistically significant depends critically on the MCMC sampling (through credible intervals). We show that using tighter criteria for significance than were used in [10] produces a more focused list of candidates for genes under positive selection. While many expected genes that have been shown to be associated with drug resistance appear on both lists, we show that the Window model often better identifies regions within these proteins that are known to be antibiotic binding sites based on X-ray crystal structures. In contrast, we show that the codon-level analysis (Independent model) has a bias toward identifying more codons under selection in large ORFs, suggesting it is more susceptible to noise. Based on these observations, we argue that the Window model in GenomegaMap is the most reliable approach for identifying genes under selection, which results from the smoothing effect achieved by modeling localized variations in w within regions of a protein (with adjustable borders). As validation, we show that, a similar list of genes under positive selection is derived from an independent set of over 5,000 Mtb clinical isolates from 15 other countries (from the CRyPTIC collection), also using the Window model.

The data on positive selection and genetic variants in both collections of clinical isolates (Moldova and CRyPTIC) is available online (https://orca1.tamu.edu/selection/) as a public resource.

## 2. METHODS

### 2.1. Sources of Genomic Data

Two sets of Mtb clinical isolates were analyzed in this paper: 2,057 isolates from a previous study in Moldova, and 5,195 globally-diverse isolates from the CRyPTIC collection. Raw sequencing data (fastq files) for isolates from Moldova were downloaded from SRA using accession numbers provided in [12]. Sequencing datasets from the CRyPTIC collection were downloaded based on accession numbers obtained from Supplemental Table 3 in [10]. While isolates could not be selected based on lineage (data not available), to promote genetic diversity, we performed stratified sampling by selecting approximately 500 isolates each from 15 different countries. TB-profiler [13] was run on the isolates to identify Mtb lineage, drug-resistance genotypes, and estimated depth of coverage (using *M. tuberculosis* H37Rv as a reference). Isolates that had < 40x coverage or were potentially mixed (with more than one sublineage represented, based on characteristic SNPs) were discarded.

### 2.2. Variant Extraction and Analysis of Positive Selection

Pilon [14] was used to map read to the H37Rv genome (NC_000962.3) and extract genetic variants (in the form of .vcf files), including SNPs and indels; Pilon also identifies low-coverage and heterogeneous sites. From the .vcf files, the nucleotide sequences of each ORF were extracted from each isolate, and aggregated by ORF (as .fasta files). Codons with low-coverage or heterogeneous nucleotides, or nonsense mutations (stop codons) were converted to gap characters (’---’), which are subsequently disregarded in the calculations by GenomegaMap. GenomegaMap was then run on each ORF using scripts for the gene-level (Constant), codon-level (Independent), and Window models (https://github.com/danny-wilson/genomegaMap).

Default parameters were generally used, as defined in the example configuration files (.xml) provided in the GenomegaMap software distribution. The parameters q and k in the Nielsen-Yang model [9] of nucleotide evolution (see Eqn 1; q corresponds to Q_ij_) are modeled with log-uniform prior distributions, with half-widths of 0.1 and 0.5. The prior distribution for w in each region is specified by a gamma distribution with parameters size=1 and shape=1. For the Window model, the w values are modeled by a piecewise continuous function. During the MCMC simulation (Metropolis-Hastings sampling), proposal updates for w values are generated using a log-uniform distribution with half-width=1. A Dirichlet-Multinomial distribution (derived in [10]) is used as a likelihood function to determine acceptance probabilities for parameter changes based on the fit of the model to the set of alleles observed in each region. Boundaries between regions are updated dynamically by reversible-jump proposals [15] from a geometric distribution with p=0.033 (1-probability that adjacent residues are in the same region), which ultimately controls the sizes of regions. The initial values for the MCMC sampling are set to q=0.17 and k=1.0, and p_j_ was set to equilibrium codon frequencies (0.01639). For the MCMC sampling for the codon-level and Window models, the trajectory length was 100,000 iterations, saving parameters every 250 steps, and eliminating the first 20% of samples as the burn-in phase (to reduce the influence of initial parameters and ensure the sampler had reached equilibrium). After the MCMC sampling, the value of w for each codon was summarized over each trajectory by extracting the mean and other statistics (like range and variance), and a 95% credible interval (95%CI) was calculated for the w value for each codon.

### 2.3. Evaluation of Models on Simulated Gene Sequences

Generation of simulated genes for comparison of GenomegaMap models under the assumption of neutral evolution was done as follows. First, a phylogenetic tree based on the 2,057 Moldova clinical isolates was constructed using RAxML [16]. Then, the lowest (or most-recent) common ancestral sequence (LCA) was computed for each ORF in the Mtb genome (excluding PPE and PE_PGRS genes, and genes known to be involved in drug resistance – this exclusion was only done for generating the simulated genes). The LCA nucleotide at each site in a given gene was derived by using Fitch’s algorithm [17] to propagate observed nucleotides from leaves (2,057 isolates) up to the root. Then, each distinct mutational event (i.e. SNP associated with a particular branch point in the tree) was randomly assigned to a new site within a new gene. This resampling process resulted in 39,564 mutations permuted to new positions among 3,446,666 sites within coding regions of 3,877 genes (with the same size distribution as in the Mtb genome), which was then used to generate the 2,057 variant sequences for each isolate for each gene.

### 2.4. Protein Structure Analysis

Atomic coordinates of proteins from X-ray crystals were downloaded from the RCSB Protein Data Bank (www.rcsb.org) and visualized in UCSF Chimera (v1.19, https://www.cgl.ucsf.edu/chimera/). The proteins were displayed as ribbon backbone models, using different colors to highlight regions predicted to be under positive selection with the sliding window model in GenomegaMap. Chimera was also used to calculate distances between amino acids, e.g. distances to active site residues.

## 3. RESULTS

We used the three models in GenomegaMap to identify and compare genes under positive selection in two collections of Mtb clinical isolates. The first was a collection of 2,236 Mtb clinical isolates from Moldova, representing all reported cases in 2018-2019 [12]. The aim of the original study was to identify transmission clusters within the region. The isolates were found to be predominantly members of lineage 2 and lineage 4, and approximately one-half of the isolates were (genotypically) drug-resistant (based on DR markers to one or more antitubercular drugs), with 32.7% MDR-TB (INH+RIF) (lineage and DR distributions are shown in Supplemental Figure S1). After downloading and preprocessing the raw sequencing data and filtering out low-quality samples, 2,057 isolates were selected for analysis.

To determine which of the genes identified as under positive selection in the Moldova dataset are generally representative, versus which might be specific to that geographic region, we carried out a similar analysis in a larger collection of 5,195 isolates from 15 other countries. These isolates were also predominantly from lineages L2 and L4, but L1 and L3 were also represented. Approximately one-third of the isolates were drug-resistant, and most of these (33.4%) being MDR (markers for both INH-R and RIF-R, based on TB-profiler). Resistance to EMB and STR was high, as well; fluoroquinolone resistance was lower, at around 10%. (Histograms of the geographic distribution, lineages, and genotypic drug resistance are shown in Supplemental Figures S2 and S3). We applied the same pipeline for genome assembly, quality filtering, and calling of genetic variants as above.

### 3.1. The Window model in GenomegaMap identifies positive selection in many genes known to be involved in drug resistance

With the Window model (where different regions within each ORF are modeled with locally-constant omega values), the MCMC simulation produces a sample of parameter combinations, including a representation of the posterior probability distribution of omega values for each codon. The mean number of windows per gene was observed to be 12.0 (evaluated on the Moldova dataset), but it scaled-up linearly with gene length: the mean window size was 28.6 amino acids. However, even though residues are grouped together in local windows which share a common w value, the w estimate for each individual amino acid can be extracted from its assigned window (which can change in each iteration of the MCMC simulation) and averaged independently.

Besides the mean value of omega for each codon, the dispersion of the sample can be used to assess confidence. In [10], the criterion used for significance was if ≥ 90% of the samples had w >1 (‘samples’ refers to combinations of parameters generated from iterations of MCMC sampling, see Methods). This corresponds to a (two-sided) 80% credible interval, which is overly generous, potentially generating many false positive codons (245 genes in the Moldova set would be significant by this criterion). Instead, we employ a more stringent criterion of ≥ 97.5% of samples having w >1, equivalent to a more standard (two-sided) 95% credible interval.

Using this criterion, 53 genes were identified as having at least one codon under positive selection in the Moldova dataset (Table 1). (dN/dS scores and statistical significance for all genes are provided in Supplemental Table T1, with indications of strong positive selection using 95% confidence criterion, and weak selection using a confidence threshold of 80%, both two-sided.) Significant genes in the PPE and PE_PGRS families were excluded from Table 1, as these genes often harbor base-call errors due to difficulty sequencing (e.g. GC-rich regions, lower coverage) or ambiguities mapping reads to these repetitive families [18, 19], and these artifactual mutations can potentially cause some of these genes to appear to be under positive selection.

**Table 1.** 53 genes with statistically significant evidence of positive selection in the Moldova collection of Mtb clinical isolates. using the Window model analysis in GenomegaMap. Asterisks indicate genes also significant in the CRyPTIC collection.

|  |  |  |  |
| --- | --- | --- | --- |
| Rv0006/*gyrA* * | Rv1286/*cysN* | Rv2258c/- | Rv2940c/*mas* |
| Rv0093c/- | Rv1288/- | Rv2388c/*hemN* | Rv3109/*moaA1* * |
| Rv0229c/- | Rv1320c/- | Rv2395/- | Rv3132c/*devS* |
| Rv0275c/- * | Rv1699/*pyrG* | Rv2407/- | Rv3299c/*atsB* |
| Rv0392c/*ndhA* | Rv1830/*resR* * | Rv2447c/*folC* * | Rv3452/*cut4* |
| Rv0667/*rpoB* * | Rv1899c/*lppD* | Rv2544/*lppB* * | Rv3695/- |
| Rv0668/*rpoC* * | Rv1901/*cinA* | Rv2652c/- * | Rv3736/- |
| Rv0758/*phoR* * | Rv1934c/*fadE17* | Rv2779c/- * | Rv3795/*embB* * |
| Rv0931c/*pknD* | Rv2043c/*pncA* * | Rv2825c/- | Rv3806c/*ubiA* * |
| Rv0969/*ctpV* | Rv2048c/*pks12* * | Rv2874/*dipZ* | Rv3822/- |
| Rv0988/- | Rv2082/- | Rv2881c/*cdsA* | Rv3828c/- |
| Rv1093/*glyA1* * | Rv2176/*pknL* | Rv2885c/- * | Rv3888c/- |
| Rv1130/*prpD* * | Rv2252/- | Rv2934/*ppsD* | Rv3911/*sigM* |
|  |  |  | Rv3919c/*gid* * |

The 53 genes under positive selection in the Moldova dataset (by the Window method) contain many genes known to be involved in drug resistance, including *rpoB* (RNA polymerase, target of rifampicin), *gyrA* (DNA gyrase, target of fluroquinolones, *gidB* (ribosome methyltransferase, affects sensitivity to streptomycin), *pncA* (pyrazinamidase, activator of pyrazinamide), and *embB* and *ubiA* (target of ethambutol). *resR* (Rv1830) also exhibits positive selection in this dataset, consistent with the findings in [1], where it was shown to be associated with general resilience to antibiotics. However, although many isolates had the S315T mutation in KatG, associated with isoniazid resistance, it was not identified by Window model (see further discussion below).

For comparison, we used the Window model in GenomegaMap to identify genes under positive selection in the larger global dataset (from CRyPTIC), with isolates from 15 other geographically-diverse countries. 173 genes were found to be under positive selection in this global dataset (excluding PPE and PE_PGRS genes) using the same criterion for significance (with at least 1 region with w>1 for at least 97.5% of the MCMC samples) (see Supplemental Table T2). 19 of the 53 significant genes in the Moldova dataset were also observed in this global collection (marked with asterisks in Table 1), representing a 36% overlap. As expected, this includes many drug-resistance-related genes, like *pncA, gid, rpoB, rpoC, gyrA, embB* and *folC*. An additional gene implicated in drug-resistance that is uniquely significant in the global collection is *aac* (Rv0262c), which is an aminoglycoside acetyltransferase that provides protection against kanamycin. Other notable genes found to be under selection in both datasets include *pks12, phoR,* and *resR*. The global positive-selection gene set also includes some additional genes that have been previously reported in other studies of positive selection in Mtb, such as *whiB6* [6].

### 3.2. Gene-level analysis of positive selection (Constant Model in GenomegaMap) fails to identify many known drug-resistance-related genes

With the gene-level analysis (Constant model), 1924 genes have w>1 in the Moldova dataset, but only 61 are significant (LB of 95%CI > 1). Two-thirds of these are PPE or PE_PGRS genes; after filtering these out, 22 genes remain that exhibit statistically significant positive selection (Supplemental Table T1). The only drug-resistance-related genes among these are *pncA, gidB,* and *ubiA*. Other DR genes like *rpoB, gyrA, embB,* and *katG* did not reach significance using this model. Many of the significant genes by the Window model have w below or only marginally above 1.0 in the gene-level model, including many of the genes implicated in drug resistance mentioned above (see Figure 1a). However, the lower bound of the 95% credible interval is less than 1.0 for almost all of the 53 genes significant by the Window model. For example, *embB* has a gene-level w of 1.19, but the 95%CI lower bound is 0.73 (not significant), and is ranked #141. Only 6 genes are significant in both models (including *pncA, gidB,* and *ubiA*). The fact that the Window model captures more drug-resistance-related genes reinforces the utility of focusing the analysis on selected regions within a gene (Window model), whereas the selection signal at the whole-gene level (Constant model) can get diluted by SNPs in other less-relevant parts of the protein.

**Figure 1.**
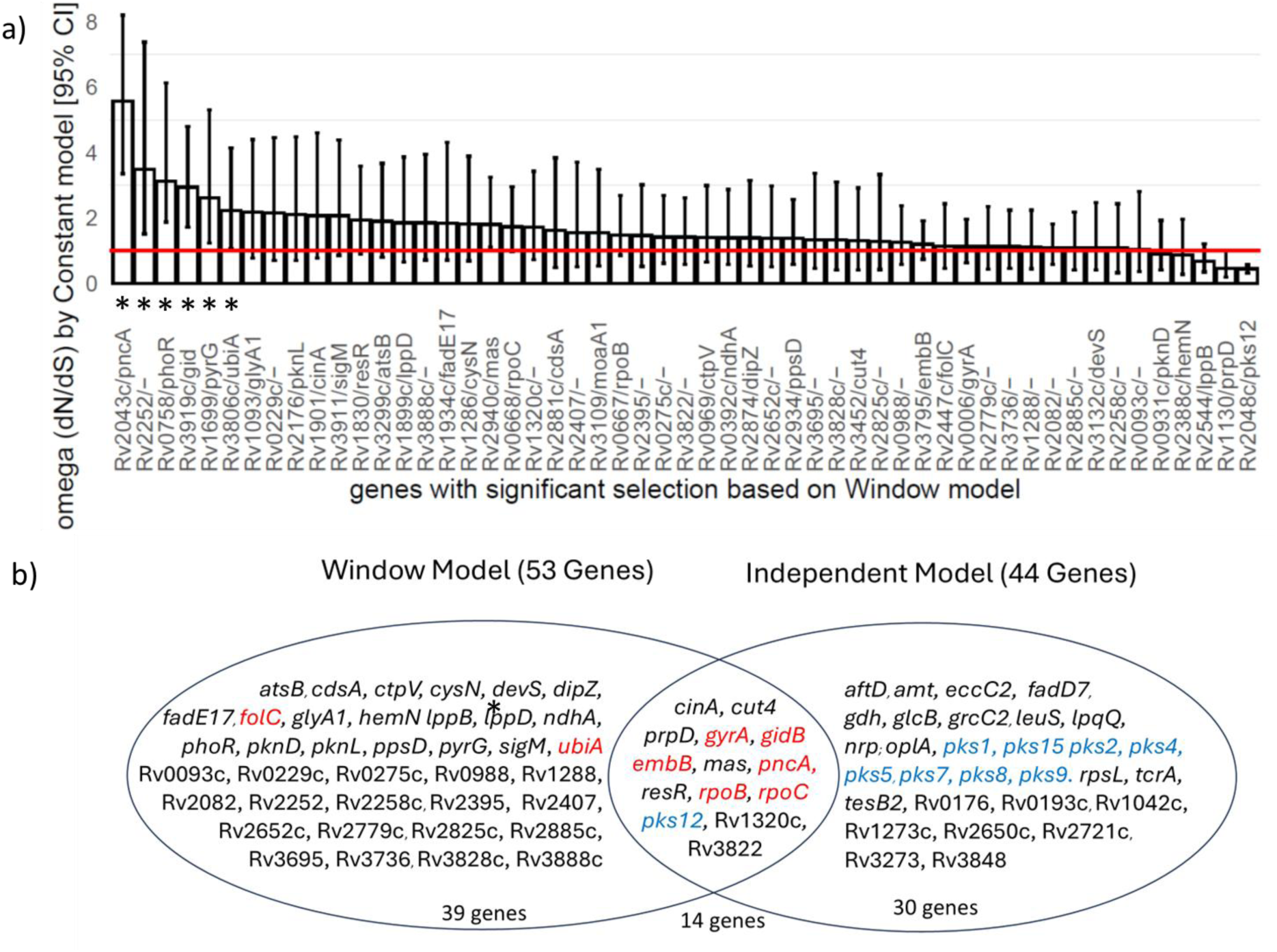
Comparison of genes under significant positive selection in Moldova dataset using 3 models. a) Barchart showing the omega estimates by the gene-level analysis (Constant model) for the 53 significant genes based on the Window model. Error bars represent 95% credible intervals for omega based on MCMC sampling. Genes that would be significant based on the Constant model (where omega lower bound>1.0, red line) are marked with an asterisk. b) Venn diagram shows overlap of 53 significant genes from Window model with 44 genes from codon-level analysis (Independent model). Drug-resistance genes are marked red; polyketide synthases are marked blue.

A similar pattern is seen for the gene-level analysis of the CRyPTIC collection (using the Constant model): 1169 genes had w>1.0, but only 42 are statistically significant (95%CI LB>1.0). Among these, only 3 genes are related to drug-resistance (*pncA, gidB, ethA*), while 4 are PPE or PE_PGRS genes (Supplemental Table T2).

### 3.3. Codon-level analysis (with the Independent model) is biased toward large ORFs

The evaluation of genes under positive selection using the codon-level (Independent) model yielded 44 significant genes in the Moldova collection, using the more stringent criterion of at least 1 codon with w>1 in ≥ 97.5% of the MCMC samples. This list of 44 genes has some overlap with the 53 significant genes by the Window model; there are 14 genes in the intersection, including most of the drug-resistance-related genes (see Venn diagram in Figure 1b). However, there are also many unique genes identified by each of the analysis methods. The Window model also highlighted *ubiA* (arabinogalactan synthesis pathway; harbors resistance mutations to ethambutol) and *folC* (resistance mutations to inhibitors of folate pathway like PAS or sulfamethoxazole), which the codon-level analysis did not.

One notable aspect of the genes exhibiting significant selection under the codon-level analysis (Independent model) is that it contains many polyketide synthase (PKS) genes. Of the 16 PKS genes in the Mtb H37Rv genome, 9 are significant in the codon-level analysis, along with *nrp* (Rv0101, non-ribosomal peptide synthase, NPRS). These modular (multi-domain) proteins are among the largest genes in the genome, with 10 out of 16 have length over 1000 amino acids (*pks12* is largest, with 4150 aa, and *nrp* is second largest, with 2511 aa). The PKS genes under positive selection are almost all unique calls for the codon-level analysis (see Figure 1b); the only PKS gene that is also detected as significant by the Window model is *pks12*. A comparison of the sizes of the genes under positive selection detected by each model is shown in the Figure 2a. Note that the size distribution of the 44 genes significant by Independent (codon-level) model is skewed higher than the 53 genes significant by the Window model, which has a distribution more similar to the overall distribution of ORF sizes in the genome (represented by the histogram on the right). Figure 2b-c shows the number of genes detected by each model in different bin sizes. When normalized by the total number of genes in each bin (Figure 2d-e), the considerable over-representation of genes in the largest bin (1200-4152 aa) by the codon-level analysis is apparent – over 40% of these largest genes contain a one or more codons under positive selection according to the Independent model (Figure 2e). By comparison, the Window model shows much less bias toward larger genes.

**Figure 2.**
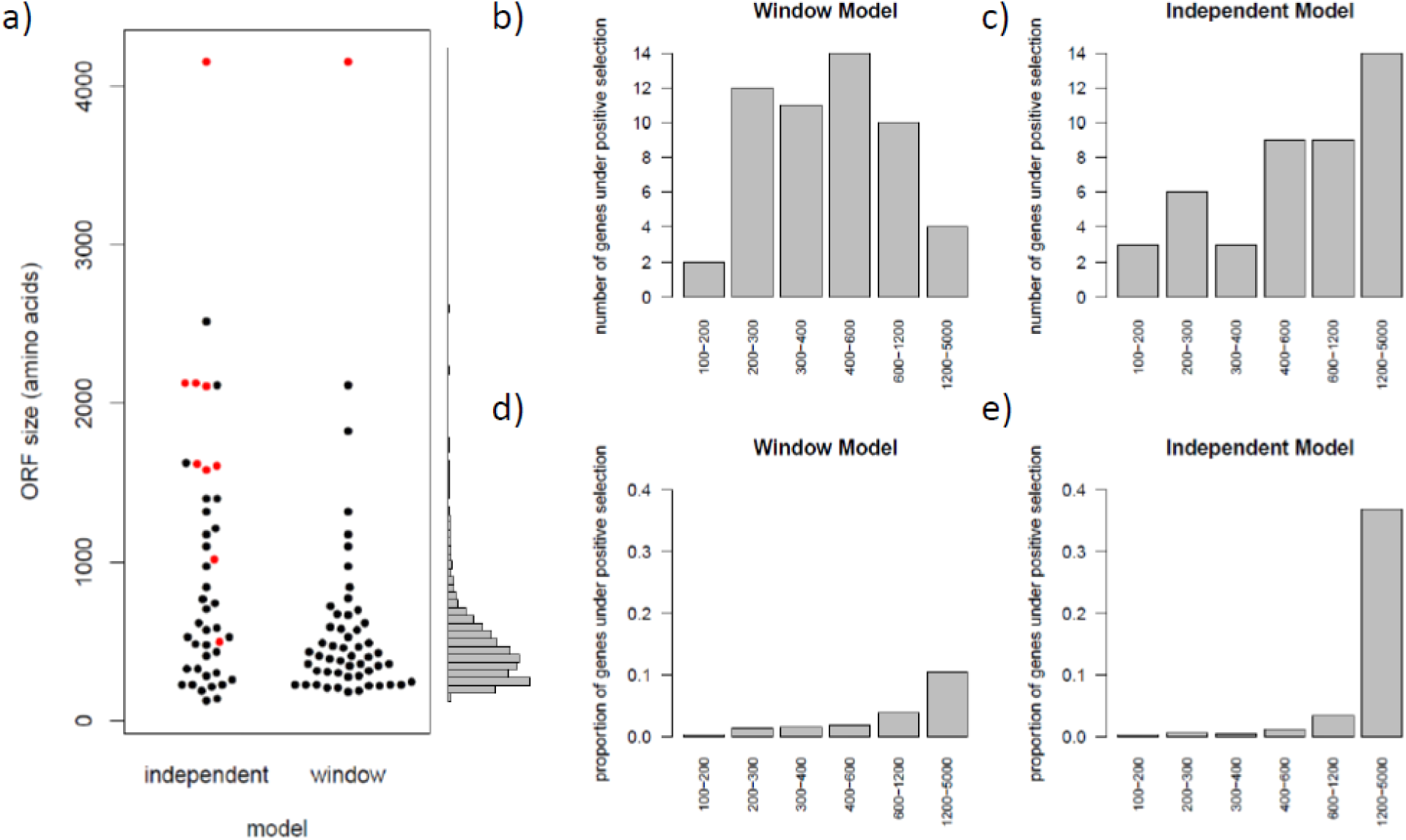
The codon-level analysis (Independent model) is biased toward larger genes. a) Beeswarm plots showing that the significant genes under positive selection by the Independent model are skewed larger than those from the Window model, whereas the Window model better matches the overall ORF size distribution (histogram to right). Polyketide synthase genes are colored red. b) and c) Number of genes under positive selection, divided into size bins (amino acids). The Independent model shows an increasing trend toward the rightmost bin (ORFs with 1200-5000 amino acids). d) and e) Proportion of genes under positive selection (counts of significant genes normalized by the overall number of genes in each size bin).

It is possible that positive selection of PKS genes could reflect biologically-meaningful selection pressures, since PKS genes make glycolipid components of the cell well that often are recognized by and module the immune response in vivo [20–22]. However, the fact that so many PKS genes show signs of positive selection could also reflect a general bias that the codon-level analysis is more likely to identify random codons as under positive selection in larger genes, which could explain why the largest genes tend to be characterized as under selection by this model.

To evaluate this hypothesis, we performed a simulation experiment to estimate the number of genes that would be expected to obtain significance by the codon-level analysis or the Window model under a Null hypothesis assuming neutral evolution. Simulated genes were generated that had mutations that occurred at the same rate as the original Moldova dataset, but evolved under the assumption of *neutral evolution*. This was achieved by first computing the lowest-common ancestral (LCA) sequence for each ORF in the Mtb genome (excluding PPE and PE_PGRS genes, and genes known to be involved in drug resistance). The ancestral sequence for each gene was copied for all isolates. Then, the sequences were mutated at random by applying mutations observed in the original genomic data. Each distinct mutational event (in any gene) (defined by a nucleotide substitution associated with a particular branch point in the original phylogenetic tree) was randomly assigned to a new site within a new gene, and the nucleotides at that site were changed to a different nucleotide for the taxa in that clade (isolates in the subtree beneath that branch point). This resampling process resulted in 39,564 distinct nucleotide changes permuted to new positions among 3,446,666 sites within coding regions of 3877 genes (with the same size distribution as in the Mtb genome). This process was used to generate a set of “random” sequences for each gene to be used for comparing the methods to evaluate positive selection, where the mutations occur at the same frequency and lineage distribution as in the original clinical isolates, but the localization of each nucleotide substitution is randomized (e.g. breaking-up clustering within ORFs) and the impact on coding (e.g. whether it is synonymous or non-synonymous) is decoupled (based on frame in new codon), hence simulating absence of selection pressure (no filtering based on translated sequence).

The 3,877 simulated gene sequences (each with variant sequences for 2,057 isolates) were then analyzed by each of the three GenomegaMap methods. The number of genes detected as significant under this Null model by the codon-level analysis (Independent model) was 66 out of 3,877 (∼1.7%), whereas only 14 of the simulated genes were identified as under significant positive selection for the Window model (∼0.4%) (see Table 2). For comparison, we also ran the gene-level analysis (Constant model) on the same set of simulated genes, which detected 16 genes with significant positive selection. Thus, the codon-level analysis would be expected to have a ∼4.5x higher false-positive rate (FPR) relative to the Window model. Furthermore, the Independent model again shows a bias toward detecting positive selection among larger genes (see Figure 3). This suggests that the codon-level analysis is more susceptible to false positive calls, due to modeling each codon with its own individual w parameter, whereas the Window model is more robust because it combines information (on genetic variants) locally over multiple codons (clusters of SNPs) (through use a constant w parameter for each segment of the ORF, with boundaries that vary over the MCMC simulation).

**Figure 3.**
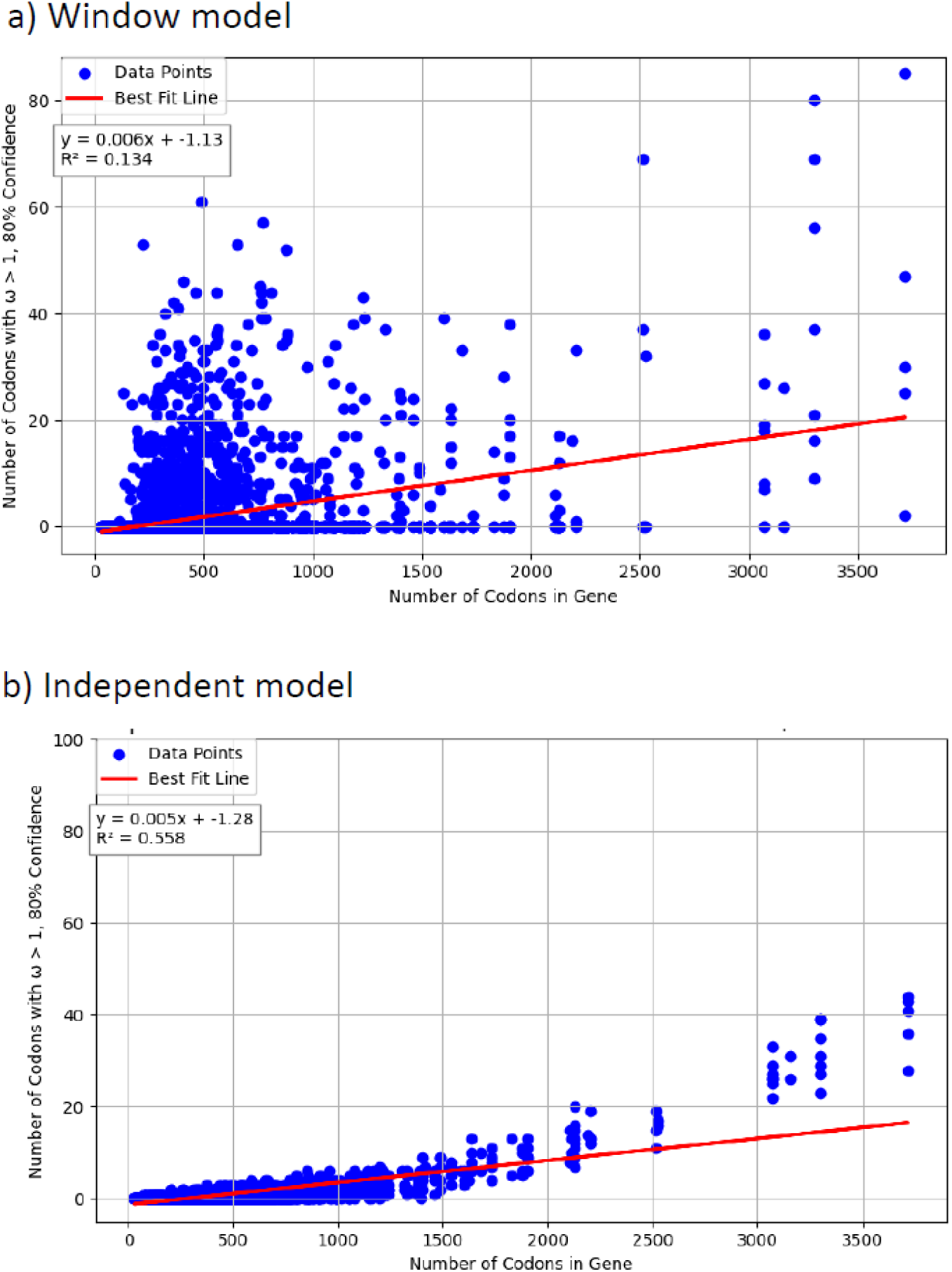
Size bias in the simulated gene set (neutral evolution). Linear regressions of the number of significant codons found against ORF among genes with weakly significant positive selection (w>1 for >90% of the MCMC samples) using a) the Window model, b) and the Independent model (codon-level analysis). The Independent model exhibits a strong correlation with ORF size (*r*^2^ = 0.558) compared to the Window model (*r*^2^ = 0.134).

**Table 2.** Evaluation of positive selection on 3877 simulated genes (under a Null hypothesis of neutral evolution).

| **GenomegaMap model** | **level of selection analysis** | **number of significant genes under positive selection (estimated FPR)** |
| --- | --- | --- |
| Constant | individual codons | 16/3877 (0.41%) |
| Window | groups of codons | 14/3877 (0.36%) |
| Independent | whole gene | 66/3877 (1.7%) |

### 3.4. The Window model identifies regions within proteins under positive selection that are likely to be functionally important

The output of the Window model (MCMC trajectory) was further analyzed to identify specific segments (or amino acids) of genes harboring the selection signal. The localization of the selection signals within each ORF can be observed in omega plots, as illustrated in Figure 4 (omega plots for *gyrA* and *embB*). These plots show the mean of the posterior distribution of omega (average of the MCMC samples) for each codon (black line). Recall that, in the Window model, even though residues are grouped in contiguous segments of the ORF which share a local w value, the window boundaries can shift between iterations of the MCMC simulation. Hence each individual residue can have its own value for the w estimate that can be extracted and averaged over the MCMC trajectory independently, which is why the mean and confidence-interval estimates of w can be plotted for each amino acid in Figure 4 and subsequent figures.

**Figure 4.**
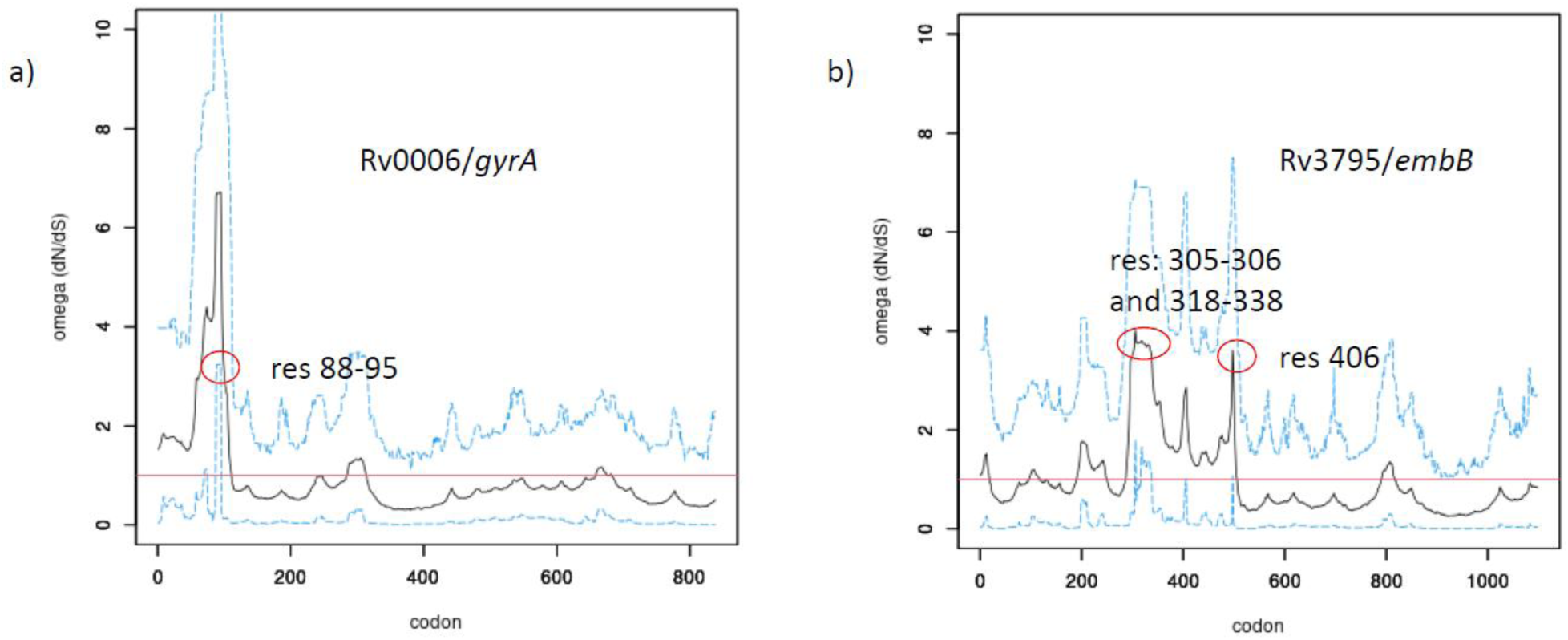
Example omega plots for *gyrA* and *embB*, showing the dN/dS for each codon estimated by the Window model. The black curve shows the mean omega value for each codon averaged over the MCMC samples; the blue lines show the upper- and lower-bounds of the 95% credible interval. Codons under significant positive selection are those where the lower blue line peaks above 1.0.

Note that the curve is smoother than the selection signal produced by the codon-level analysis (Independent model). (Omega plots for the corresponding genes based on the Independent model are shown in the Supplemental Figure S4). This smoothing is a consequence of how the omega estimate for one codon effectively combines local information on genetic variants at surrounding codons. Therefore, peaks in the w signal using the Window model tend to center on clusters of SNPs where there is a local over-abundance of non-synonymous mutations. The upper and lower bounds of the 95% CI are shown as blue dashed lines in the figure. The significant codons are (only) those for which the lower bound of the 95%CI exceeds 1.0, or when w>1 in 97.5% of the MCMC samples. In contrast, the omega plot of a gene using the codon-level model is much more variable (spiky), because the omega values at one codon is completely independent from the others around it, and there is no sharing of information locally.

In the case of GyrA, there are two significant peaks. The predominant peak consists of residues 89 to 95 (where omega peaks at 6.7). This contains the two codons most frequently associated with fluoroquinolone (FQ) resistance – A90 and D94 [23]. In fact, although FQ resistance was only ∼10% in this population (10.3% for moxifloxacin), A90V was observed in 40 isolates (1.9%) in the Moldova collection, along with D89G/N and D94G/A/N/Y/H. There is also a minor peak at residues 70-74, which is just barely significant (lower bound of 95%CI just barely exceeds 1.0).

This is in an adjacent alpha-helix, also in the DNA-binding pocket; A74S been reported in FQ-resistant clinical isolates, but is less frequently observed [24, 25]. Although, some compensatory mutations in *gyrB* have also been reported to be associated with FQ-resistance [26], none of the codons in *gyrB* were statistically significant using the Window model.

Regions associated with resistance mutations were also found to be significant in *embB* (Figure 4b), which is in the arabinogalactan (AG) synthesis pathway. The three codons most commonly observed to be mutated in ethambutol-resistant mutants– residues 306, 406, and 497 [27] – all exhibit a significant w value in the Window model. In addition, residues 305 (adjacent to 306) and 317-338 (in between 306 and 406) are highlighted as significant. While no mutation in residue 305 in EmbB has yet been associated with resistance, some sites in the 317-338 window have (such as 319 and 328; see WHO resistance mutation catalog, [28]). In addition, in *ubiA*, residues 171-176 have significant w values. UbiA is also in the AG pathway, and these residues have been reported in EMB-resistant mutants [29].

In the CRyPTIC collection, *aac* (Rv0262c, aminoglycoside acetyltransferase) was identified to be under positive selection. Codons under positive selection an *aac* span 86-94. These residues form a loop that binds CoA (rather than the aminoglycoside), based on crystal structure of the complex of *aac* with kanamycin (PDB: 1M4I; see Supplemental Figure S5).

It is notable that *pncA* and *gidB*, two of the genes with the strongest mean selection signals (mean w=2.93 and 1.94, respectively), show selection in regions throughout the gene (151 and 78 codons respectively, see Supplemental Figure S6), which is consistent with the diversity of resistance mutations observed, including loss-of-function [30]. *pncA* is an activator of pyrazinamide, and *gid* is a ribosomal rRNA methyltransferase that natively facilitates binding of streptomycin [31, 32]. Hence any mutations that abrogate function of these two genes could potentially confer resistance [33]. This could explain why GenomegaMap detects a significant signal for positive selection that is diffused across both of these targets.

#### 3.4.1. RpoB

Rifampicin resistance mutations are commonly observed in of *rpoB* (beta’ chain of RNA polymerase), specifically in the Rifampicin Resistance Determining Region (RRDR), residues 435-452, including the most frequent mutation S450L in RIF-R mutants), which defines the physical binding site for rifampicin. As observed in the omega plot, there are two strong peaks indicating regions of the ORF with significant selection (see Figure 5a). The first peak (residues 435-451), with omega values exceeding 4, clearly corresponds to the RRDR. The second peak consists of residues 488-496. These have been reported to be associated with rifampicin-resistant clinical isolates in China [34]. Since they co-occurred with mutations in the RRDR region of *rpoB*, it was speculated that they could be compensatory. We observe this in the Moldova collection too. In fact, based on the phylogenetic reconstruction (generated using RAxML), there are two clear cases where mutation events in the 488-496 region occurred in lower branches after mutations in the RRDR. For example, 15 isolates with the *rpoB*:I488V were embedded in a cluster of 111 isolates with *rpoB*:S450L (see Supplemental Figure S7). In crystal structures of the RNA polymerase, the residues 488-496 are in a parallel beta strand to 426-452 in the RIF-binding pocket (with Ile491 directly adjacent to Ser450; see Figure 5b).

**Figure 5.**
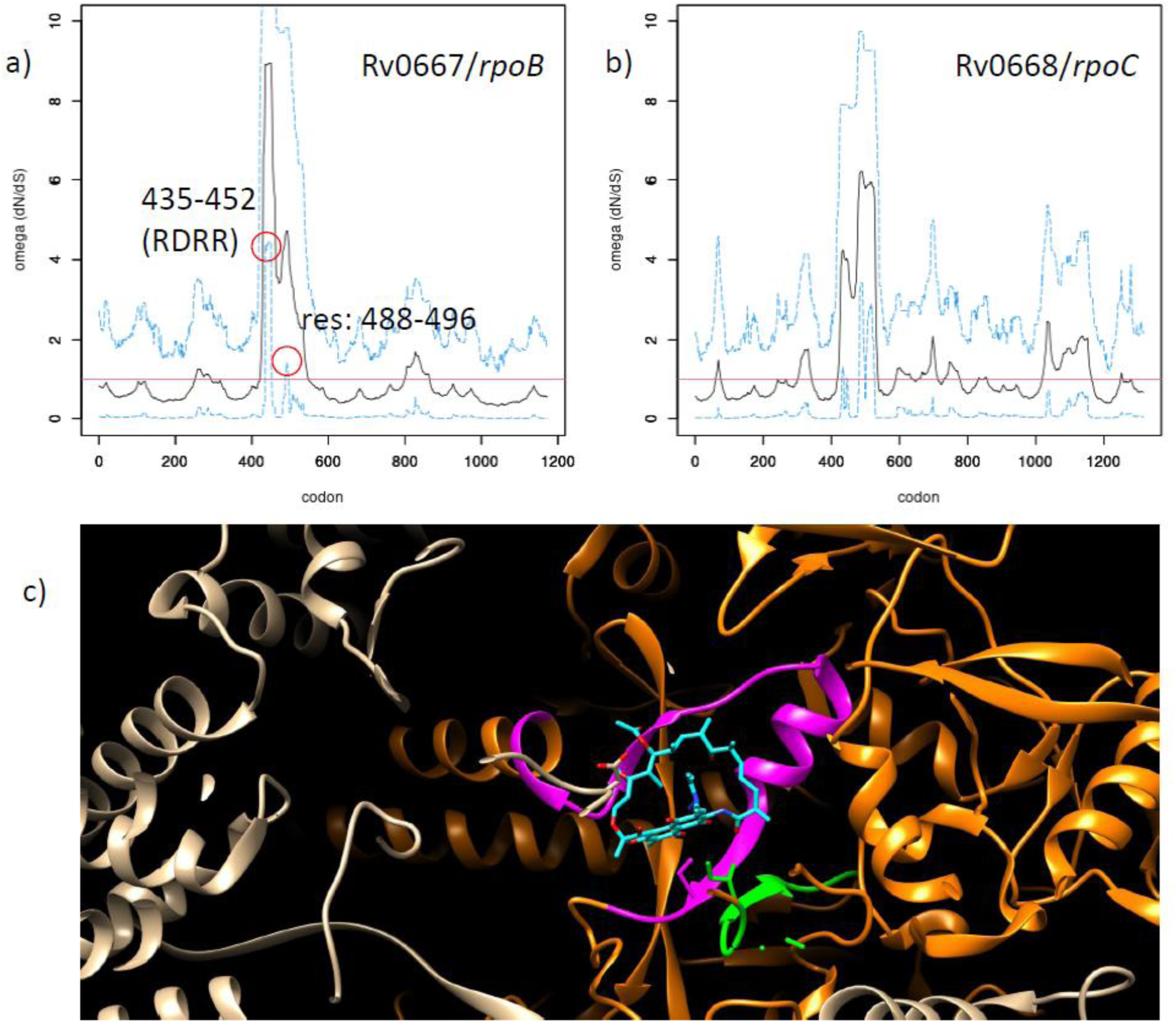
Analysis of positive selection in RpoB and RpoC. a) Omega plot for *rpoB*. b) omega plot for *rpoC*. c) X-ray crystal structure of complex of RNA polymerase (*rpoABC*) (PDB: 5uhc) with rifampicin (cyan), showing that 488-496 (green) forms a parallel beta-strand to 426-452 (purple); both are regions under positive selection.

In addition to RIF-resistance mutations in *rpoB*, compensatory mutations have been observed in *rpoC*, especially around residues 432-527 [7]. Indeed, *rpoC* is also in the list of 53 genes exhibiting positive selection in the Moldova clinical isolates (using the Window model). As shown in Figure 5c, several regions of codons attain significance, including 431-436, 445-447, 482-498, and 502-527, which corresponds well with the region containing compensatory mutations described by Comas et al [7].

#### 3.4.2. CinA

CinA was found to be under significant positive selection in the Moldova dataset, and has recently been implicated in INH tolerance [35]. The activity influencing drug sensitivity was localized to the N-terminal pyrophosphatase domain of the two-domain protein. The omega plot (Figure 6a) exhibits a significant peak for residues in the 82-86 range (in the pyrophosphatase domain). Genetic variants observed in this region include: T82I, V83F, and V86A. These residues are in an alpha-helix proximal to Asp80, which projects into the active site and coordinates Mg^2+^ bound to the diphosphate in NADH, in a crystal structure of CinA homolog from *T. thermophilus* (PDB: 4uux) (see Figure 6b). Mutating Asp80 (D80A) abrogates enzyme activity and increases INH sensitivity [35]. In fact, one of the isolates in the Moldova collection had D80E, which likely also affects CinA function. However, none of the isolates with these mutations in CinA were genotypically resistant to INH, ETH, DLM, or PTM (based on drug-resistance markers analyzed by TB-profiler). In contrast, only residue 421 was detected as under positive selection in CinA by the codon-level analysis.

**Figure 6.**
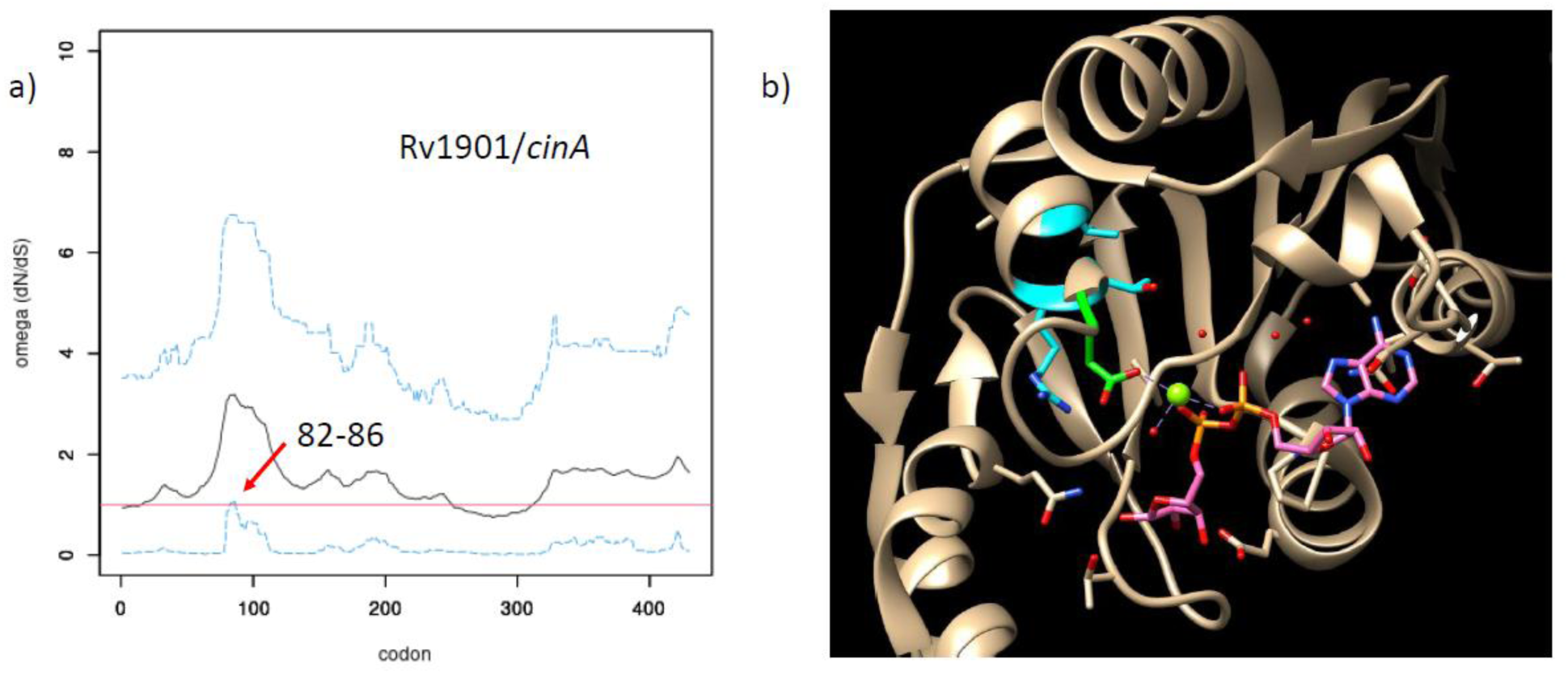
Analysis positive selection in CinA. a) Omega plot for *cinA*, showing selection of residue 82-86 (where lower blue line peaks above 1.0). b) Crystal structure of CinA from *T. thermophilus* (N-terminal domain) (PDB: 4uux); green: Asp80 (proximal to NADH in active site, purple), blue residues: codons under positive selection.

#### 3.4.3. PhoR

*phoR,* which has been found to be under positive selection in previous studies of Mtb clinical isolates [36], has among the strongest signals for positive selection in both the Moldova dataset and the CRyPTIC dataset. In the Moldova dataset, the codons in *phoR* with genetic variants under positive selection include: 52-84 and 145-159 (in the periplasmic sensor domain) and 188-201 (in the HAMP domain; an alpha-helical domain that facilitates oligomerization by form coiled-coils, often utilized in signal transduction proteins) (see Figure 7).

**Figure 7.**
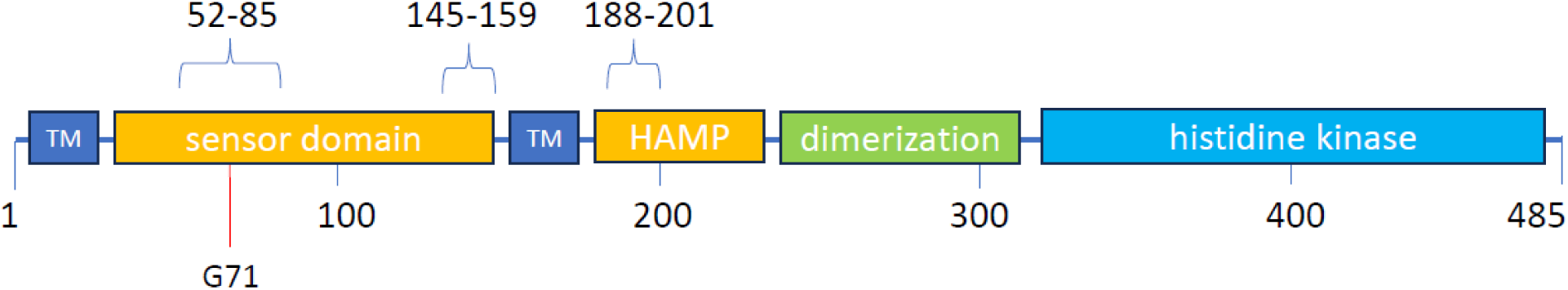
Regions under positive selection in *phoR*, mapped onto domain structure. ‘TM’ refers to transmembrane domains. The residue (G71) that differentiates the *Mtb* and *M. bovis* orthologs is indicated.

### 3.5. ESX-1 genes exhibit signs of positive selection

Several components of the ESX-1 complex (a type VII secretion system that is important for virulence [37, 38]) exhibit positive selection in the larger global collection of Mtb clinical isolates, including *eccCa1, eccD1*, and *eccE1*, along with *espA*. In addition, *eccB1, esxA* (ESAT-6)*, and espFIJ* all contained segments showing evidence of positive selection by the weaker definition of significance (>90% of MCMC samples had w>1). Only *espB* and *espK* showed weak significance in the Moldova dataset, possibly because of the smaller number of isolates. The region of *espA* under positive selection includes codons 67-72 (Figure 8a). Non-synonymous genetic variants in this region include: K66R/Q, N67T, R68A, N69S, H70Y, V71L, N72K. These residues lie in an alpha-helix of a predicted 4-helix bundle, proximal to the WXG motif (res. 55-57) (see Figure 8b), which forms a loop that plays a critical role in recognition by the ESX-1 pore complex [39]. As noted above, *whiB6* and *phoR,* which both regulate expression of ESX-1 [40, 41], also exhibit positive selection in the global dataset.

**Figure 8.**
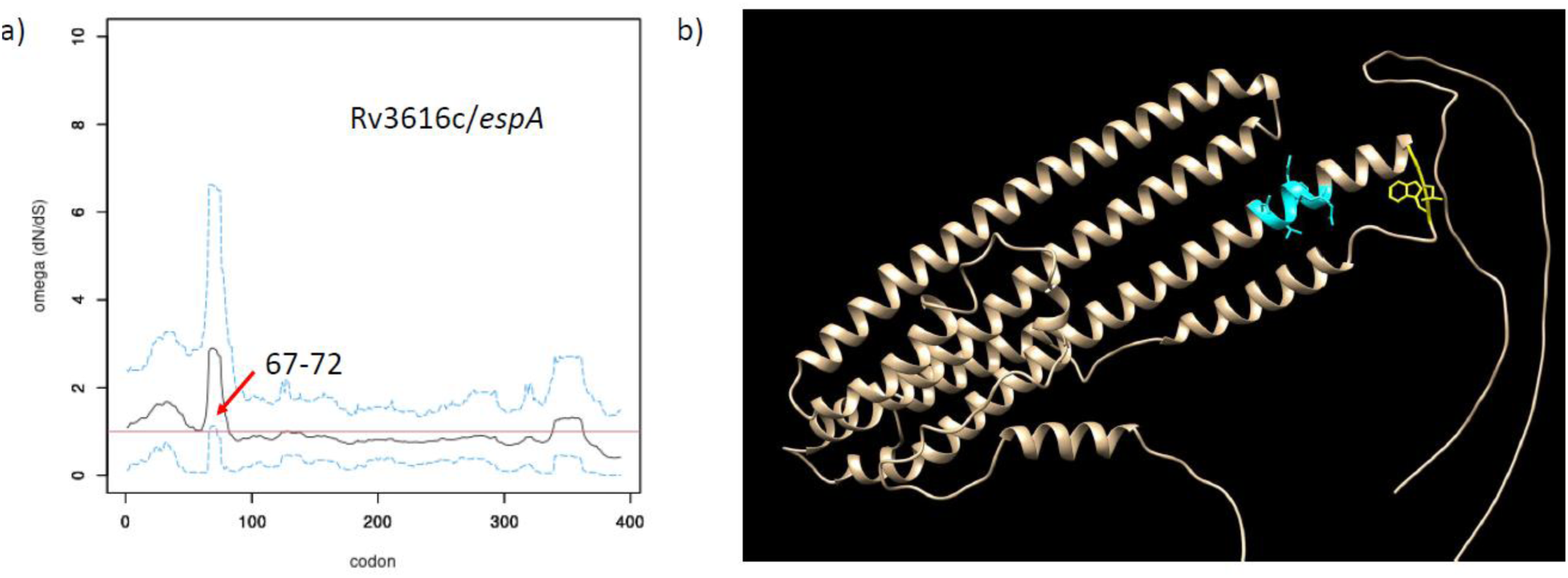
Analysis of positive selection for EspA. a) Omega plot for *espA,* showing significant selection for codons 67-72. b) AlphaFold model of EspA. Residues under positive selection (67-72) are colored blue. The WXG motif (residues 55-57), involved in recognition by the ESX-1 pore complex, are colored blue.

## 4. DISCUSSION

Mtb is considered well-adapted to its lifecycle as an obligate human pathogen, and the genome is fairly homogeneous (low diversity) [42]. Strong evidence for purifying selection has been found throughout the genome [43]. Nevertheless, sporadic SNPs and other polymorphisms are often observed in genomes of individual isolates (presumably arising within patients during the course of infection [44]), which are infrequent (unfixed alleles, appearing in only one or a few isolates), reflecting on-going evolution [45]. Some of these mutations might reflect adaptations to pressures related to modern clinical treatment of TB, including chemotherapy, but possibly also effects of co-morbidities (e.g. diabetes, COPD, malnutrition), co-infection (e.g. HIV), which could involve exposure to non-TB antibiotics as well as impacting patient immune status, oxidative environment, etc. Statistical methods for evaluating positive selection are a powerful tool for identifying genes adaptive to these pressures in large genomic databases.

There have been multiple studies investigating positive selection among genomes of Mtb clinical isolates, based on principles ranging from phenotype-genotype associations [6], to SNP clustering (e.g. nucleotide diversity, mutation burden) [1, 4], to homoplasy [4], convergent evolution [5], traditional dN/dS analysis [3, 13], and phylodynamics [2]. In addition to genes already known to be involved in drug-resistance, these studies have identified other genes which may play an adaptive role to modern clinical stresses, such as *resR* [1] and *phoR* [36].

Most of these studies evaluated selection at the gene-level. However, selection can be analyzed at other scales, including codon-level and clusters of SNPs (intermediate level). The purpose of this study was to evaluate and compare the results of all three levels of analysis.

One of the potential problems with gene-level analysis is that the selection signal, which might derive from one or a few residues (e.g. in an active site) could get diluted from mutations in other less-relevant parts of the ORF, when aggregating SNPs over the whole ORF. On the other hand, analyzing each codon independently cannot take advantage of clustering of SNPs, which often occurs in active sites, or sites of protein-protein interaction. We used GenomegaMap as a tool, which implements all 3 analyses within a Bayesian framework, and uses MCMC sampling to make inferences about parameters (omega) and their certainty (credible intervals).

We found that the intermediate (or “Window” model) provides the most compelling list of candidate genes. The Window model segments the ORF into multiple contiguous regions, and allows each region to have a different w parameter, but assumes the codons within a local region all share the same omega value. Many of the genes found to be under positive selection by the Window model are known to be involved in drug resistance, (*rpoB, gyrA, embB, pncA, gidB, ubiA, folC*…) which would be expected to be under positive selection in a collection of Mtb clinical isolates from patients receiving chemotherapy, especially in a geographic region with high incidence of drug-resistance (such as Moldova [12]). In addition, on the list of candidates from the Window model, we observed other genes previously reported as targets of positive selection in other studies, such as *resR* [1] and *phoR* [36]. *prpD* is a methylcitrate dehydratase (propionate metabolism pathway) that is regulated by *prpR*, which was identified as associated with multi-drug tolerance [46]. Rv0988 and Rv2082 have both been reported in another study of positive selection in clinical isolates resistant to bedaquiline and clofazimine [47]. While the function of Rv2082 is unknown, Rv0988 is an ABC-transporter implicated in preventing acidification and phagosome maturation [48]. This consistency for “expected” genes under positive selection with previous studies increases confidence that other candidate genes (including those in other pathways which might not be directly related to drug-resistance) might also be genuinely under positive selection and playing an adaptive role.

Another advantage of the Window model of selection analysis is that it can identify specific regions (or residues) within an ORF that are under positive selection. In the case of genes commonly associated with drug resistance, the regions under selection were often found to co-occur with known drug-binding regions of the protein. This co-location of selection signals with active sites or structural binding sites for antibiotics lends confidence that the Window model is detecting genuine signals of selective pressure. As implemented in GenomegaMap, this model assumes w is constant locally but varies among regions. However, the borders between regions also get adjusted dynamically during the MCMC sampling, producing a smoothly varying profile of ω across the ORF. We define significant codons to be those where >97.% of the sampled w values from the MCMC sample (the lower-bound of the 95% credible interval) exceed 1.0, which is more stringent that the criterion used in the original paper [10]. In cases where protein crystal structures are available, the regions under positive selection often correspond well to parts of the protein involved in interacting with inhibitors (often in active sites), such as in *rpoB* (residues in the RDRR that bind rifampicin) and *gyrA* (residues binding fluoroquinolones), as well as compensatory regions that might reduce the fitness cost of resistance mutations.

Another compelling example is the proximity of the residues under selection in *cinA* to the active site of the pyrophosphatase domain, and in *aac*. The insights about which parts of a protein are under selection are helpful in understanding the mechanisms driving it, beyond just labeling a gene as being under positive selection. The correspondence of codons under selection to structural information (e.g. for known drug-binding sites) lends additional confidence in the list of candidates identified by the Window model.

Individual codons under selection can also be highlighted by the Independent model in GenomegaMap (where each codon has its own w parameter). However, we observed that many of the genes on the list of candidates from the codon-level analysis were among the largest genes in the genome, and this size bias suggests the Independent model might be more susceptible to making type I errors. In fact, we used a simulated set of genes with sequences evolved under the assumption of neutral evolution (with the same distribution of ORF sizes and nucleotide content as the clinical isolates from Moldova, but with observed phylogenetic changes permuted randomly to new sites) to show that the codon-level analysis (Independent model) has a roughly 5 times higher estimated false positive rate than the Window model. This is likely due to how the Independent model treats each codon independently, making it more susceptible to errors from random coincidences of SNPs, whereas the Window model is more robust because of how it integrates information over multiple codons in a region (by forcing the codons in a region to share the same w), and hence takes advantage of clusters of SNPs where there is a local excess of non-synonymous over synonymous SNPs.

Based on the analysis of *rpoB* with the Window model, we observed signals of positive selection in the RDRR region (residues 435-452), as well as in an another segment (residues 488-496) that, which, while sequentially discontiguous, packs against the RDRR in 3D space, forming part of the same RIF-binding pocket. This could provide a structural explanation for how mutations at one site could mitigate the fitness cost of mutations at the other. The fact that several cases were observed where the mutations in 488-496 region of *rpoB* came after the mutations in the RDRR (as subsequent evolutionary events) supports the hypothesis that mutations in the 488-496 region might be adaptive and help reduce the fitness cost of the earlier mutations in the RRDR. We also observed *rpoC* to be under positive selection, which is is consistent with the finding of compensatory mutations for RIF resistance in this gene [7].

Among drug-resistance-related genes that were *not* detected as significant by the Window model were *katG* and *inhA*, which is unexpected, given the high level of (genotypic) INH resistance in the region (>40%). None of the three models identified *katG* (activator of isoniazid) as positively selected in this collection, despite mutations in *katG* being the most frequently observed in INH-resistant isolates. Wilson [10] also noted that *katG* was not found to be under positive selection by the Window model in their larger dataset, but was identified by the codon-level analysis (Independent model) because the majority of INH-resistance mutations occur in a single codon - Ser315. However, this could also be easily identified by complementary tools such as homoplasy analysis [4], as mutations at this site often occur multiple times independently in clinical populations. *inhA* was probably not detected because mutations occur predominantly in the promoter region upstream of *fabG1*, more frequently than in the coding region, affecting expression levels of *inhA*).

Several genes potentially involved *indirectly* in INH resistance were observed to be under positive selection, including CinA and NdhA. CinA is an NADH-metabolizing enzyme whose biological role is not well-understood [35]. Though CinA is non-essential under regular growth conditions (7H9 medium), it was found to be conditionally more essential (decreased Tn insertion counts) in the presence of isoniazid (INH) [49]. It was subsequently shown to be involved in NADH recycling, and to mitigate the impact of INH-NADH adducts [35]. It was also shown to affect MICs for delamanid (DLM) and pretomanid (PTM), which also generate NADH adducts. CinA has two domains: an N-terminal pyrophosphatase and C-terminal deamidase. The signal for positive selection was associated with the pyrophosphatase domain, and the genetic variants observed (in the Moldova collection) where proximal to the active site. The proximity of these mutations to the active site suggest that CinA is under genuine selective pressure in this population, the source of which is still unknown. Another gene observed to be under positive selection that is potentially related to INH resistance indirectly was was *ndhA*. *ndhA* (Rv0392c) is one of two type II NADH dehydrogenases, along with *ndh* (Rv1854c) which can compensate for INH-NADH adduct formation by shifting the intracellular NADH/NAD+ ratio [50]. Although, historically, *ndh* is more often observed to acquire resistance mutations [51], mutations in *ndhA* have also been reported to be associated with INH resistance [52].

On the list of 53 genes identified to be under positive selection in the Moldova dataset, we observed several regulatory proteins, including *pknD* and *sigL* (ser-thr protein kinases), *sigM* (sigma factor), Rv1320 (adenylate cyclase), *phoR* and *devS* (two-component sensors).. These regulators could be involved in adaptations to various stresses. For example, *devS* plays a critical role in regulating the response to hypoxia (low oxygen tension, e.g. inside granulomas), a hallmark of the host immune response to TB infection [53–56], and the intracellular oxidative environment could also be influenced by antibiotics (such as isoniazid, which generates oxidative radicals in the process of activation [57]). *whiB6* (under positive selection in the CRyPTIC collecion) is a transcriptional regulator that was found to be associated with multi-drug-resistance in a GWAS study of clinical isolates from Peru [6]. *whiB6* has also been implicated in expression of ESX-1 [40]. Several metabolic genes also show signals of positive selection in this population (*cysN* - sulfate adenyltransferase/kinase; *glyA1* – serine hydroxymethyltransferase; *hemN* - coproporphyrinogen III oxidase; *pyrG* - CTP synthase, *atsD* - arylsulfatase; *fadE17* - acyl-CoA dehydrogenase; *cdsA* - CDP-diglyceride pyrophosphorylase).

Metabolic shifts often accompany changes in response to stress [58], though the specific role of these pathways remains to be elucidated. *phoR* had among the strongest signals for positive selection in both datasets (Moldova, and global collection). There is strong evidence for *phoR* being under positive selection in clinical isolates in previous studies [36]. *phoR* is part of a two-component sensor, PhoPR [59], which plays a key role in virulence [60, 61]. *phoR* is membrane-bound sensor coupled to a histidine kinase that phosphorylates *phoP*, which binds DNA and acts as transcription factor. PhoPR function is required for virulence, as disruption mutants are highly attenuated in mice [60, 61]. A mutation in *phoP* is one of the few genetic differences between H37Rv and H37Ra, to which the difference in virulence of these strains has been partially attributed [62]. As a part of a two-component sensor, *phoR* regulates many lipid synthesis and cell-wall biogenesis genes [41].

PhoPR has been implicated in responses to several types of environmental stress, including acidic pH [60]. Using phylodynamics, Chiner-Oms et al. [36] found that *phoR* exhibited an increase in selection of non-synonymous variants, compared to other animal-adapted members of the MTBC (Mycobacterium Tuberculosis Complex), that coincided roughly with expansion of the pathogen into the human population. This and other evidence suggests mutations in *phoR* might facilitate transmission among humans and might have been a key step in the evolution of Mtb into a human pathogen [63]. The lineage-specific polymorphism that differentiates Mtb *phoR* from the ortholog in *M. bovis*, which is associated with decreased transmission in humans, is G71I [63], which falls in the middle of the sensor domain. Gonzalo-Asensio et al. showed that recombineering this mutation into L2 and L4 strains actually down-regulates expression of the PhoPR regulon. Using the Window model we observed naturally-occurring genetic variants (non-synonymous mutations) in both the sensor domain and HAMP domains of PhoR. The non-synonymous mutations in the sensor domain might cause similar functional changes that are ultimately beneficial and lead to selection in Mtb clinical isolates. Mutations in the HAMP domain will have to be further investigated to determine their impact on PhoPR function and understand how this can lead to adaptation.

In order to determine whether the genes under selection might be specific to Moldova, we applied the same analysis of positive selection to a larger collection of Mtb clinical isolates from 15 other countries. Many of the drug-resistance genes, along with genes such as *resR* and *phoR* were also detected as significant in the global dataset. There were approximately 3 times as many genes (173) with statistically significant positive selection as in the Moldova dataset (53). This is possibly due in part to the increased incidence of unfixed mutations (often unique SNPs observed in only one or a few isolates), the probability of which increases with size of the collection (number of genomes, barring duplications due to outbreak clusters), or possibly due to the increased diversity among isolates selected from 15 different countries. However, in theory, dN/dS analysis should factor-out differences in overall mutation rate, and instead identify genes where there is a *relative* excess in the rate of non-synonymous over synonymous SNPs, which is not necessarily a consequence of aggregating SNPs from a larger collection of genomes.

Among the genes under positive selection in the larger global collection, we observed several genes from the ESX-1 cluster. In particular, *eccCa1, eccD1, eccE1* were under positive selection, along with *espA*. ESX-1 is one of five type-VII secretion systems (T7Ss) in Mtb [37]. It plays an important role in interacting with the host immune system, including phagosome-maturation arrest, host cell membrane lysis, inhibition of autophagy [64], and up-regulation of host-cell genes and cytokine signaling [65–67], and is required for virulence in animal models of infection [38]. ESX-1 secretes a number of effectors (including *esxA* and *esxB*, along with other *esx* and PPE proteins), and can disrupt host-cell membranes and modulate cytokine expression [67, 68]. The ESX-1 locus (Rv3864-Rv3883c) contains members of the transmembrane pore complex (*eccA1-E1*), the *mycP1* protease, and several secreted effectors (*espBEFGHIJKL, esxAB, pe35, ppe68*). Based on analogy to a cryo-EM structure of the ESX-5 complex from *M. xenopi* [69], *eccCa1, eccD1,* and *eccE1* are predicted to interact and form the cytoplasmic face of the transmembrane pore. Interestingly *espACD* are in a separate locus (Rv3514c-Rv3616c) from the main ESX-1 cluster. Specifically, *espA* binds to the ESX-1 pore complex and facilitates secretion of EsxA and EsxB [39, 65]. Thus, we speculate that the genetic variants observed in ESX-1 genes could play a role in fine-tuning the host immune response. ESX components have been identified in other studies of positive selection. For example, *esxW* (secreted via ESX-5) was identified to harbor convergent mutations in different Mtb lineages in a collection of isolates from Viet Nam, which was proposed to help explain the increased transmission of the Beijing strain in the geographic region [70]. *esxW* was not under significant positive selection in our global collection, and the only *esx* protein that was under significant positive selection was *esxH*, a member of ESX-3. ESX-1 proteins *eccD1* and *eccE1* were also observed to be under positive selection in the analysis of over 51,000 clinical isolates by Liu et al. [1].

Increased selection of genetic variants in the ESX-1 system could be the result of adaptation to modern TB treatment paradigms (e.g. shortening of disease course through chemotherapy, better clinical control of inflammation) that might benefit from adjustments in host-pathogen interactions (e.g. enhancement or suppression of immune response). It is interesting to note that PhoPR has been shown to regulate expression of ESX-1 and *esxA* (ESAT-6) secretion [71]. Similarly, *whiB6*, which is also on the list of genes under significant positive selection in the global collection, also regulates expression of ESX-1 (demonstrated in *M. marinum*; [40]). The selection signals for these two regulators lend additional support to the potential adaptive role of genetic variants affecting ESX-1 function in this global sample of clinical isolates.

## 5. LIMITATIONS

One of the limitations of GenomegaMap is that it does not take into account phylogenetic information when analyzing residues for positive selection, and hence is not sensitive to mutations that are selected multiple times independently (homoplasy). This was done for efficiency, to avoid have to build phylogenetic trees, which does not scale-up well for thousands of genomes. While most regions under positive selection exhibit a cluster of SNPs enriched for non-synonymous over synonymous mutations, there are some cases where mutations can be focused primarily on a single site, such as for the KatG:S315T mutation, which GenomegaMap had trouble detecting. However, GenomegaMap can easily be complemented with other tools, such as for homoplasy analysis (e.g. SNPPar [72]). Pepperell [4] and others have demonstrated that homoplasy analysis can effectively identify residues and genes under positive selection, including those associated with drug resistance among Mtb clinical isolates.

A related limitation is that GenomegaMap is restricted to identifying positive selection in coding regions, because of its reliance on interpreting mutations in terms of non-synonymous versus synonymous substitutions (i.e. to evaluate dN/dS). However, methods like homoplasy analysis and SNP burden tests that focus on evolution or clustering (and do not rely on translation) can again be used to complement GenomegaMap to identify signs of positive selection in non-coding (or intergenic) regions. For example, Liu et al [1] used a SNP burden test to assess positive selection in promoter regions upstream of ORFs, suggesting that the mutations might produce their phenotypes through altering gene expression levels.

While we found that the Window model of GenomegaMap worked better than the gene-level or codon-level analysis for identifying regions under selection, we did not systematically explore the effect of window size. Using default parameters, amino acids were grouped into windows of approximately 28 amino acids. This produced a smoothing effect on omega values (averaged over the MCMC trajectory) because it allowed neighboring residues to share the same local omega parameter (depending on the location of boundaries). It is possible that by varying the parameter *p* of the geometric distribution in the MCMC simulation, average window sizes could be increased or decreased, which could have influenced the sensitivity for detecting regions under positive selection (by over- or under-smoothing). For example, it could be that enforcing smaller window sizes might allow increased resolution by focusing more precisely on clusters of residues sporting an excess of non-synonymous mutations, undiluted by mutations in surrounding residues.

The implementation of the Window model in GenomegaMap implicitly assumes that residues under positive selection are clustered in sequentially contiguous regions (segments of an ORF). However, it is possible that discontiguous residues fold together in 3D space to be functional, such as in some active sites. Actually, these are sometimes recognized as *separate* windows under positive selection, as we illustrated with the case of *rpoB*, though finding a way to combine them could increase statistical sensitivity, rather than analyzing the regions independently. A variety of extensions to the method could be imagined, such as higher-level clustering of residues with elevated ω estimates, but the challenge would be revising the Bayesian model to calculate the statistical significance for these clusters in a rigorous way.

Another limitation of GenomegaMap is that it does not take into account indels (insertions/deletions), only nucleotide substitutions. However, indels can play a critical role in activating/deactivating proteins, in some cases producing phenotypes. For example, Indels are commonly observed as a mechanism of resistance in non-essential drug-activator genes, like *pncA* [30], and have also been reported in genes related to PDIM synthesis (*ppsA-E, mas*), which have been found to produce a slight growth advantage in laboratory settings [73]. There is a 7-bp insertion that divides *pks15/*1 into two ORFs and eliminates the synthesis of PGL (phenolglycolipid) found in more virulent strains of Mtb (e.g. W-Beijing clade in lineage 2) [21]. Some indels (including large-scale deletions spanning >10kb and tens of genes) can serve as lineage-specific markers [74, 75]. Indels have recently been found to be a driver of variation among PPE and PE_PGRS genes (relying on long-read sequencing for accuracy) [76]. Association of indels with specific phenotypes, such as drug resistance, can be evaluated through statistical methods such GWAS (Genome-Wide Association testing), when aligned and encoded properly as a genotype [77].

A final limitation is that GenomegaMap does not work as robustly on genes in the PPE and PE_PGRS families because difficulty sequencing these regions [18] can generate incorrect or uncertain base calls, which can inflate the apparent mutation rate in such genes. This can be related to higher GC-content, which causes problem for Illumina sequencing technologies and often leads to lower coverage and increased base-call ambiguity [19]. We observed that many PPE and PGRS genes were identified as being under positive selection by GenomegaMap, though some of these might be artifacts (false positives). These genes are mostly secreted, structural proteins (non-enzymes) [78], and almost all are non-essential in-vitro (e.g. knockout by transposon mutagenesis has no identifiable phenotype, [79]). Few genes in these families have been found to have identifiable functions, with a few notable exceptions, such as PPE51 [80, 81]. However, some of these genes have been implicated in pathogenesis via modulation of host immune cells [82, 83], for example, PGRS33 [84]. In fact, several PPE and PGRS genes have been highlighted in another study of positive selection (also based on dN/dS) among Mtb clinical isolates in China [3]. Therefore, we provide selection data (from GenomegaMap) on these genes in the Supplemental Tables for completeness.

## 6. CONCLUSION

In summary, GenomegaMap is an effective tool for evaluating genes under positive selection that scales up well to analyzing large genomic databases. Furthermore, our results suggest that evaluating selection at an intermediate scale (using the “Window” model) identifies the most credible list of candidate genes under selection, compared to gene-level analysis (the Constant model) and codon-level analysis (the Independent model). The Window model takes advantage of clusters of SNPs with a local increased rate of non-synonymous over synonymous mutations, which amplifies signals of selection over the other two models. Some of the genes under selection are likely responding to antibiotic treatment (anti-TB chemotherapy), whereas other genes exhibiting positive selection might be reflecting adaptations to other unknown pressures related to the clinical environment and represent avenues for further investigation into TB biology and potential new drug targets.

## Supplemental Data

**Supplemental Table T1**. Selection scores and statistical significance for all ORFs in the Mtb H37Rv genome for the Moldova dataset (2,057 clinical isolates). Results for 3 models in GenomegaMap are provided: gene-level analysis (Constant model, one omega parameter per gene), Window model (omega values shared by codons locally within regions), and codon-level analysis (Independent model, separate omega parameter for each codon). For the Window and codon-level models, the number of significant codons (“strongly selected codons”) are given, using a 95% credible interval as a significance criterion (w>1 for >97.5% of the MCMC samples). (The number of weakly selected codons using 80%CI is given too.) The genes are sorted by the “max_omega_LB” which is the peak height of the lower blue curve in the omega plots (lower bound of the 95%CI). The 53 significant genes by the Window model (excluding PPE/PGRS genes) are marked in yellow.

**Supplemental Table T2.** Selection scores and statistical significance for all ORFs in the Mtb H37Rv genome for the global Mtb genome collection (5,195 clinical isolates from 15 countries). Results for the Window model in GenomegaMap are provided: The number of significant codons (“strongly selected codons”) are given, using a 95% credible interval as a significance criterion (w>1 for >97.5% of the MCMC samples). (The number of weakly selected codons using 80%CI is given too.) The genes are sorted by the “max_omega_LB” which is the peak height of the lower blue curve in the omega plots (lower bound of the 95%CI). The 178 significant genes by the Window model (excluding PPE/PGRS genes) are marked in yellow.

## Acknowledgements

Supercomputing time was provided by the High Performance Research Computing (HPRC) center at Texas A&M University.

## CRediT authorship contribution statement

TRI - Conceptualization, Data Curation, Formal Analysis, Writing - original manuscript; AS - Visualization, Writing – review and editing.

---

**Supplemental Figure S1.**
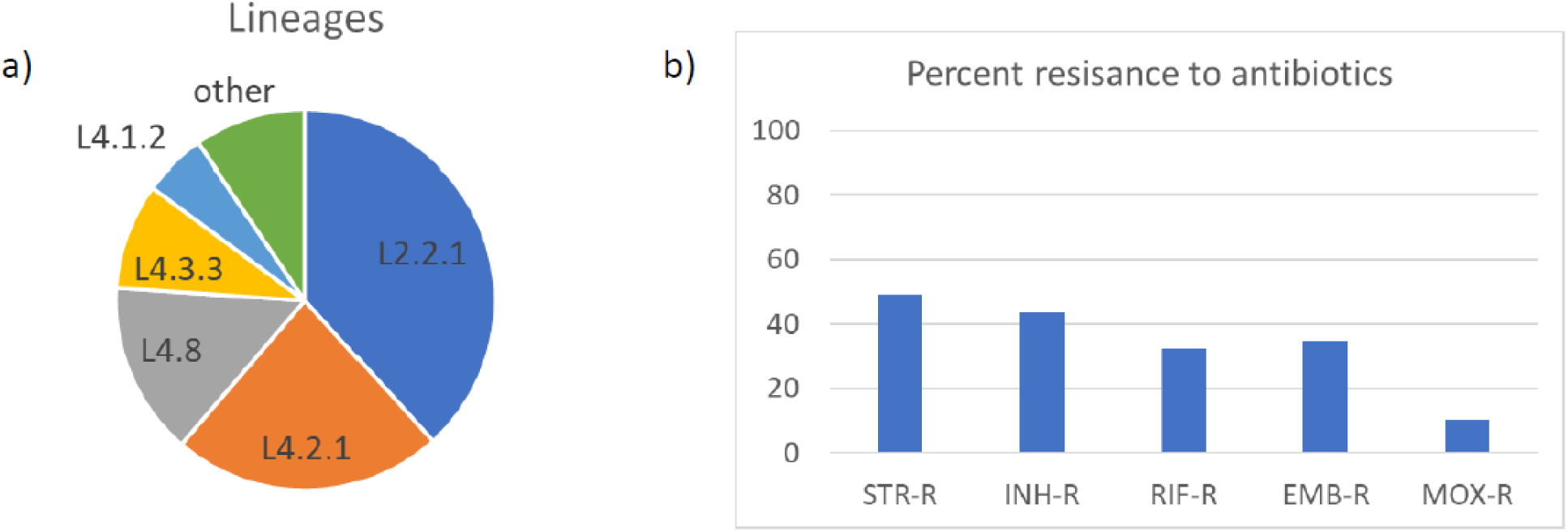
Lineage distribution and drug-resistance profile for 2,057 Mtb clinical isolates from Moldova (as determined by TB-profiler).

**Supplemental Figure S2.**
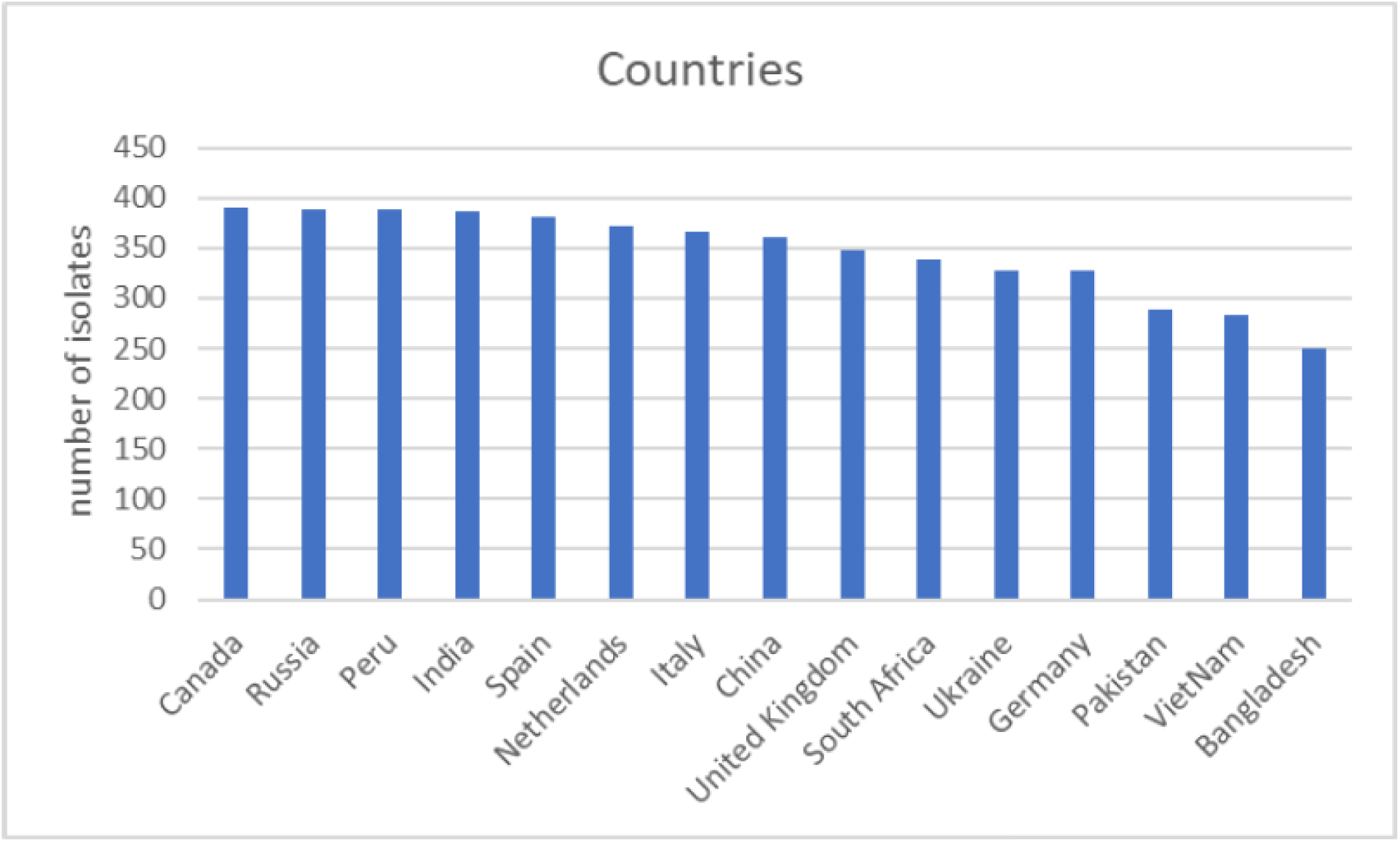
Countries represented in global collection of 5,195 Mtb clinical isolates (from CRyPTIC collection).

**Supplemental Figure S3.**
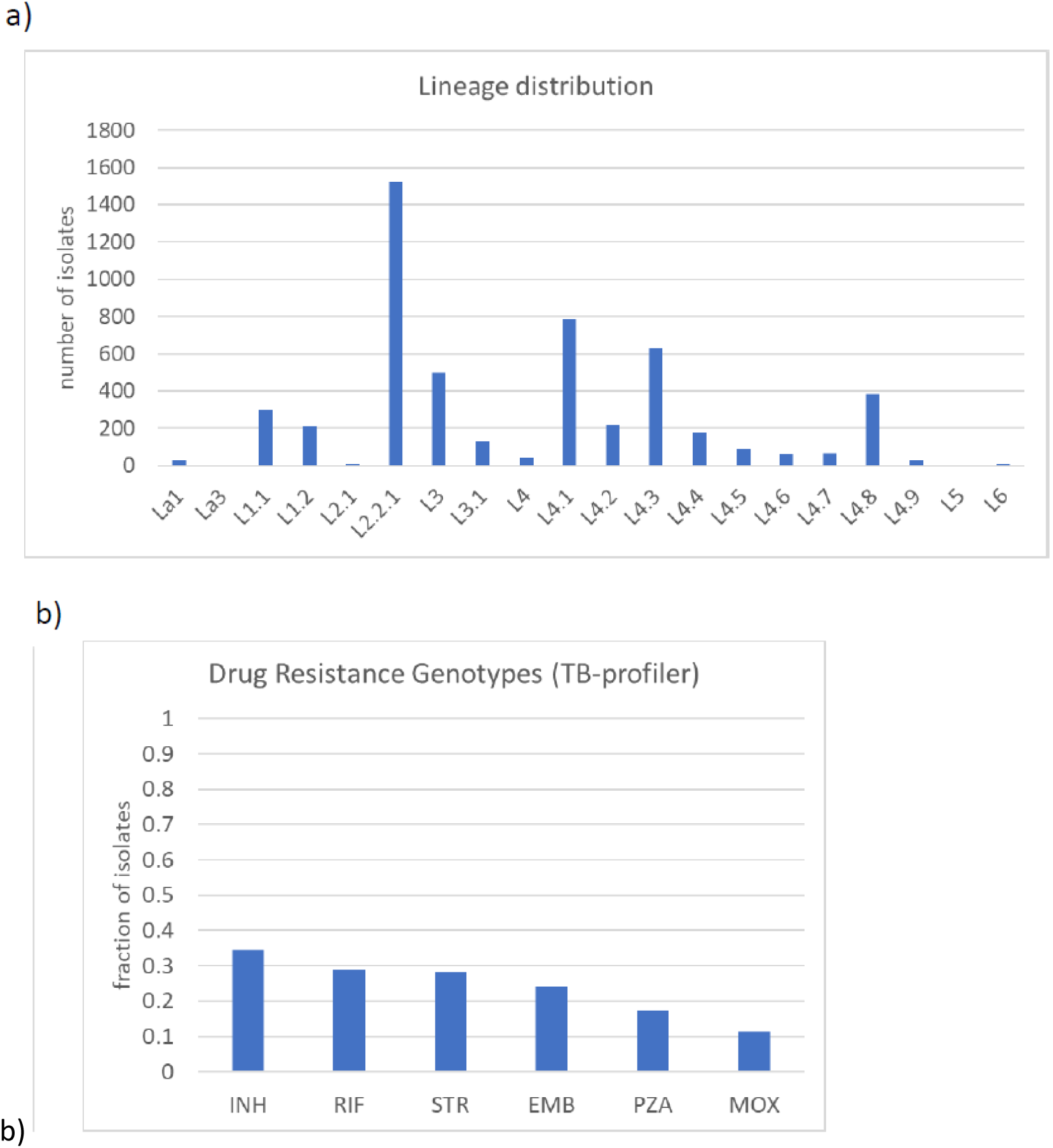
Lineages and drug-resistance genotypes represented in global collection of 5,195 Mtb clinical isolates.

**Supplemental Figure S4.**
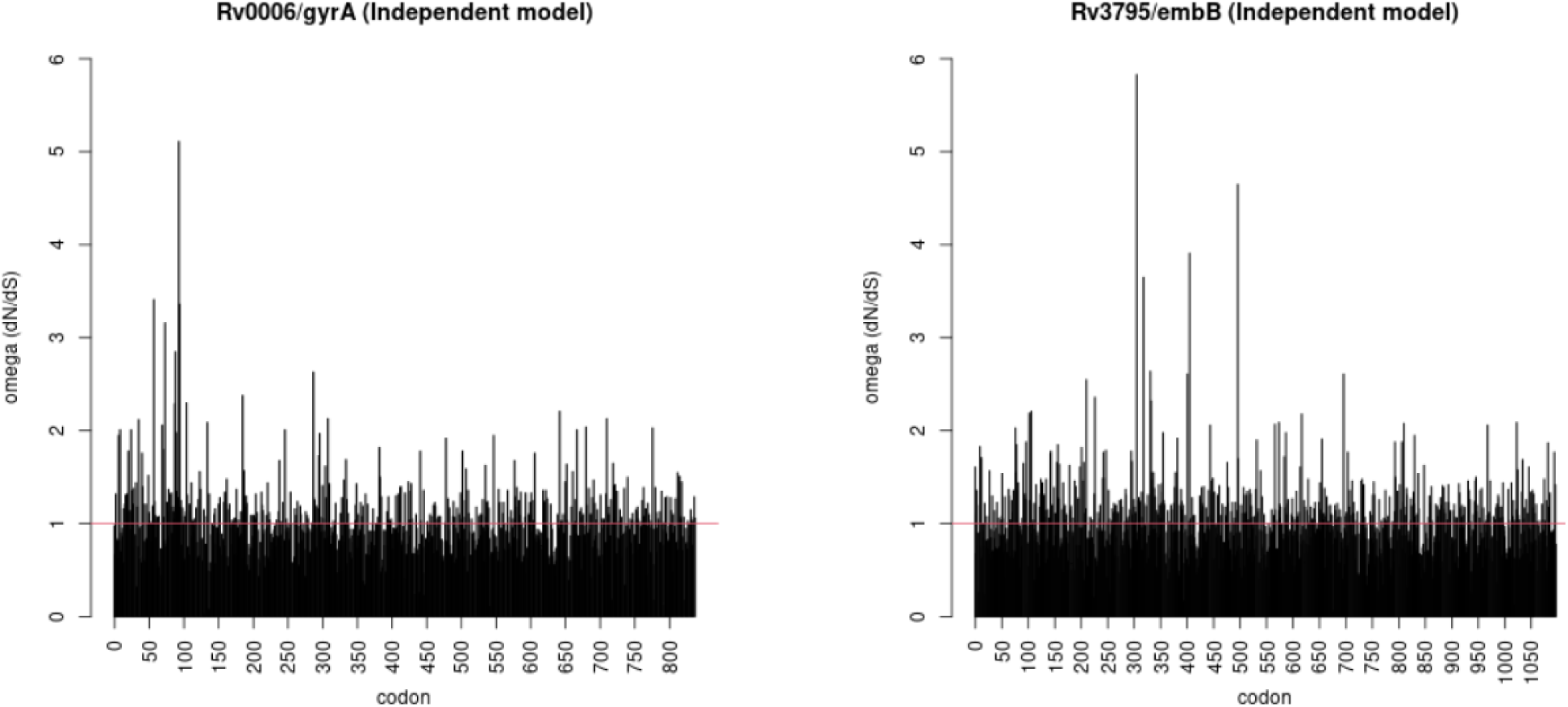
Omega plots for *gyrA* and *embB* based on the Independent model (codon-level analysis).

**Supplemental Figure S5.**
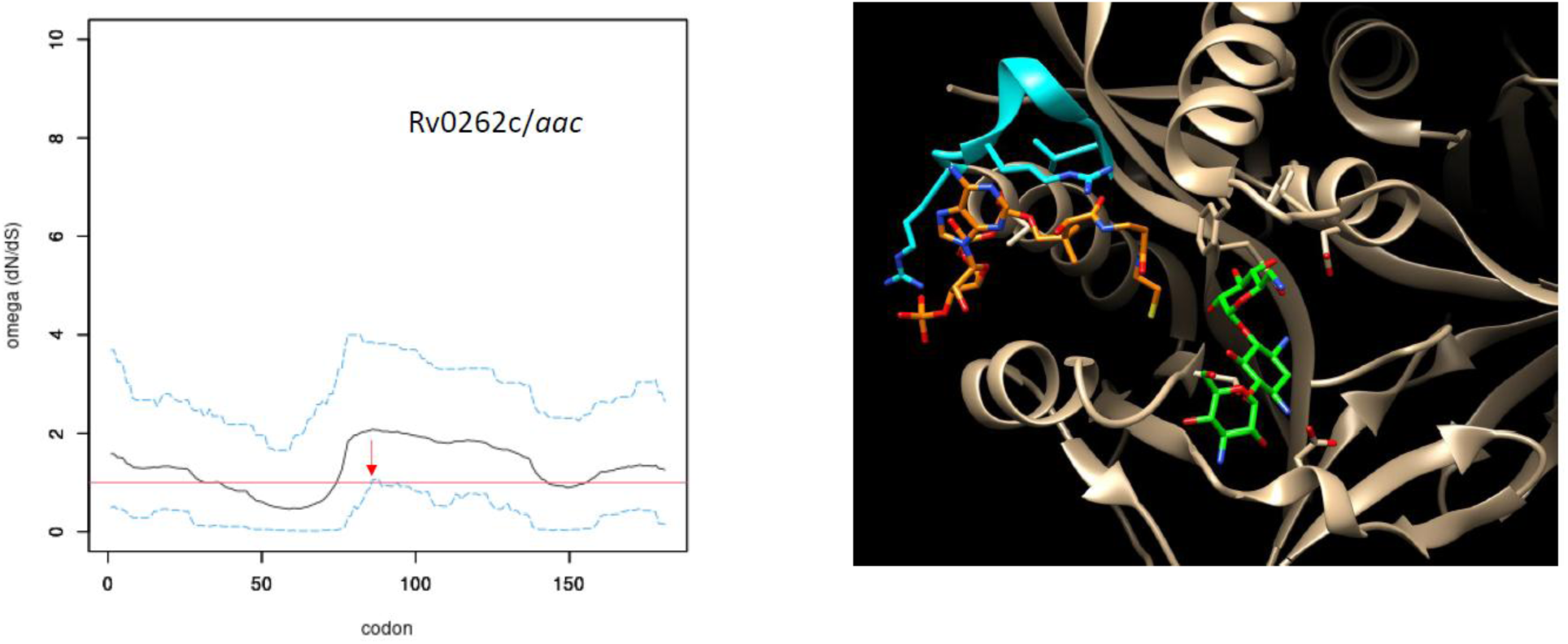
Analysis of positive selection for *aac* (Rv0262c, aminoglycoside 2’-N-acetyltransferase). a) Omega plot for *aac*: b) Genetic variants in *aac* mapped onto crystal structure (PDB: 1M4I) of complex of Mtb *aac* with CoA (orange) and kanamycin (green); residues under selection (86-94) are marked in blue.

**Supplemental Figure S6.**
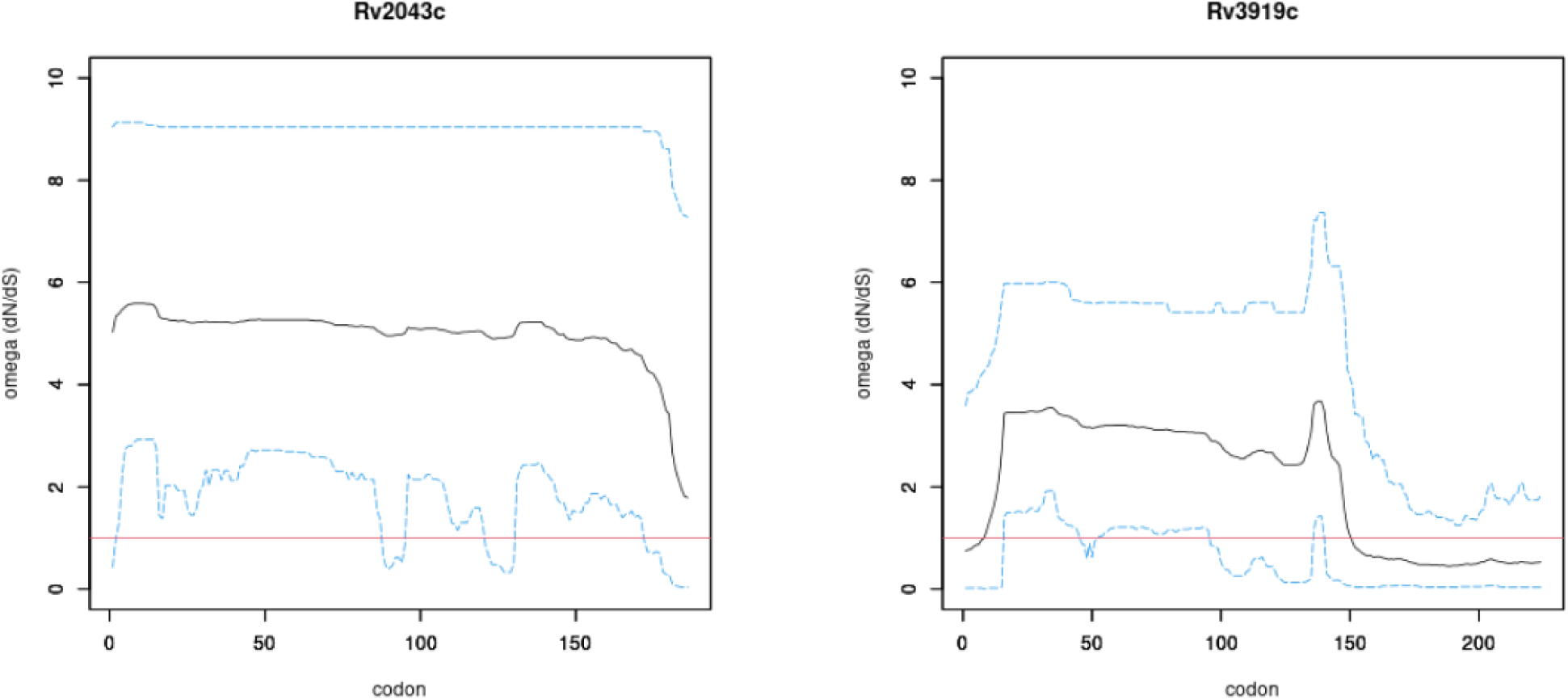
Omega plots for *pncA* (Rv2043c) and *gidB* (Rv3919c) based on the Window model, showing a high fraction of the coding regions under positive selection (lower blue line > 1).

**Supplemental Figure S7.**
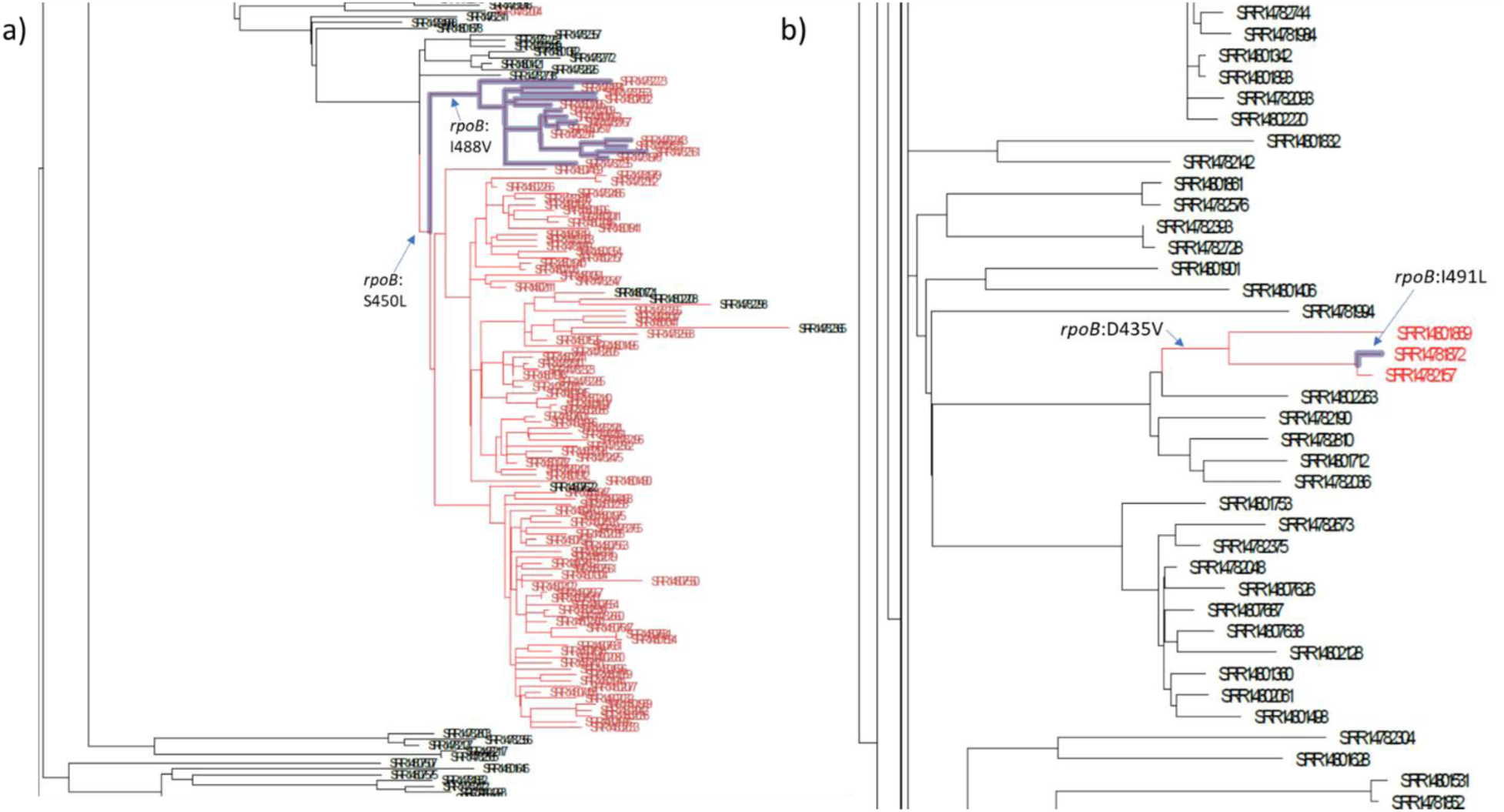
Portions of the phylogeny of Moldova isolates showing the order of acquisition of RpoB mutations (the leaves are SRA accession numbers of isolates). a) The *rpoB*:I488V mutation in the highlighted clade of 15 isolates occurred after the *rpoB*:S450L mutation (111 isolates, taxa highlighted red), within the L2.2.1 lineage. b) The *rpoB*:I491L mutation in a single isolate (highlighted branch) occurred after the *rpoB*:D435V mutation within a clade of 3 isolates (red) in the L4.3.3 lineage.

